# Essential role of hyperacetylated microtubules in innate immunity escape orchestrated by the EBV-encoded BHRF1 protein

**DOI:** 10.1101/2021.06.11.448012

**Authors:** Damien Glon, Géraldine Vilmen, Daniel Perdiz, Eva Hernandez, Guillaume Beauclair, Frédérique Quignon, Clarisse Berlioz-Torrent, Vincent Maréchal, Christian Poüs, Marion Lussignol, Audrey Esclatine

## Abstract

Innate immunity constitutes the first line of defense against viruses, in which mitochondria play an important role in the induction of the interferon (IFN) response. BHRF1, a multifunctional viral protein expressed during Epstein-Barr virus reactivation, modulates mitochondrial dynamics and disrupts the IFN signaling pathway. Mitochondria are mobile organelles that move through the cytoplasm thanks to the cytoskeleton and in particular the microtubule (MT) network. MTs undergo various post-translational modifications, among them tubulin acetylation. In this study, we demonstrated that BHRF1 induces MT hyperacetylation to escape innate immunity. Indeed, the expression of BHRF1 induces the clustering of shortened mitochondria next to the nucleus. This “mito-aggresome” is organized around the centrosome and its formation is MT-dependent. We also observed that the α-tubulin acetyltransferase ATAT1 interacts with BHRF1. Using ATAT1 knockdown or a non-acetylatable α-tubulin mutant, we demonstrated that this hyperacetylation is necessary for the mito-aggresome formation. Similar results were observed during EBV reactivation. We investigated the mechanism leading to the clustering of mitochondria, and we identified dyneins as motors that are required for mitochondrial clustering. Finally, we demonstrated that BHRF1 needs MT hyperacetylation to block the induction of the IFN response. Moreover, the loss of MT hyperacetylation blocks the localization of autophagosomes close to the mito-aggresome, impeding BHRF1 to initiate mitophagy, which is essential to inhibiting the signaling pathway. Therefore, our results reveal the role of the MT network, and its acetylation level, in the induction of a pro-viral mitophagy.

## Introduction

The innate immune system provides the first line of defense against different invading pathogens. This process is based on the sensing of motifs from the foreign organism, called the pathogen-associated molecular pattern (PAMP), by different host pattern recognition receptors (PRR). In the case of a viral infection, their recognition notably leads to the induction of the interferon (IFN) response, which induces the expression of interferon-stimulated genes and the synthesis of cytokines known for their antiviral properties (Lee and Ashkar, 2018).

Mitochondria carry out a crucial role in many cellular processes ranging from energy production to programmed cell death, from calcium homeostasis to cell immunity. They constitute a platform for signaling pathways involved in innate immunity thanks to the mitochondrial-resident protein MAVS (mitochondrial antiviral signaling protein), predominantly localized at the mitochondrial outer membrane surface (Arnoult *et al*., 2011; West *et al*., 2011). RIG-I (retinoic acid-inducible gene) and MDA5 (melanoma differentiation-associated protein 5), two cytoplasmic PRRs that detect viral genomes, are notably translocated to the mitochondria to interact with MAVS that recruits and activates TBK1 (TANK-binding kinase 1). This kinase is required for the phosphorylation of the transcription factors IRF3 and IRF7 (interferon regulatory factors 3 and 7), leading to the subsequent activation of type I IFN promoter (Arnoult *et al*., 2011; West *et al*., 2011). The functions of mitochondria depend on their morphology, which is in turn dependent upon mitochondrial dynamics, including fission, fusion, and motility. Moreover, mitochondria are actively recruited to specific cellular locations to respond to their functions. In eukaryotic cells, the cytoskeleton and notably microtubules (MTs) play a critical role in the distribution of mitochondria throughout the cytoplasm by facilitating their transport to areas with high metabolic demands (Frederick and Shaw, 2007; Kruppa and Buss, 2021).

The MT cytoskeleton is a highly dynamic polymer of α and β-tubulin heterodimers, which is involved in a variety of cellular functions, such as supporting cell structures, maintaining cell polarity, but also allowing the transport of organelles. Moreover, it has been shown that the MT network of virally infected cells could be used for intracellular transport of viral particle/genomic material to the sites of replication or assembly (Zheng *et al*., 2017). A broad range of MT-associated proteins (MAPs) contributes to regulating MT dynamic behavior. MAPs include kinesins and dyneins, which both allow the positioning of organelles and the long-distance transport of vesicles along MT tracks. Kinesins generally transport their cargos toward the plus-end of MTs at the cell periphery. Conversely, dyneins transport their cargos to the minus-end of MTs in the cell center (Hancock, 2014). The relationship between MTs and mitochondrial transport is not completely understood. Mitochondrial movement involves mitochondrial adaptors such as small mitochondrial Rho GTPases Miro1/2 and TRAK1/2 (López-Doménech *et al*., 2018). Indeed, Miro proteins can engage both kinesin and dynein to mediate the bidirectional movement of mitochondria along MT tracts (Fransson *et al*., 2006; Frederick and Shaw, 2007; Tang, 2018).

MT functions are regulated by numerous post-translational modifications such as acetylation, detyrosination, phosphorylation, methylation or polyglutamylation (Janke and Magiera, 2020). Among them, the acetylation of α-tubulin, which mainly occurs on the Lysine in position 40 (K40), has emerged as an important regulator of numerous cell functions (Perdiz *et al*., 2011). For example, α-tubulin acetylation, which is often linked to microtubule stabilization, modulates the binding of MAPs (Dompierre *et al*., 2007; Mackeh *et al*., 2014; Reed *et al*., 2006) and therefore may modulate organelle movement. Indeed, stable MTs are particularly involved in mitochondrial transport (Friedman *et al*.,2010), and it has been reported that acetylation of α-tubulin mediates dynein-dependent transport of mitochondria along MTs (Misawa *et al*., 2013).

Our group recently unravelled the novel functions in innate immunity of the Epstein-Barr virus (EBV) BHRF1 protein, a viral Bcl-2 homolog (Khanim *et al*., 1997), related to its mitochondrial localization (Vilmen *et al*., 2020). We demonstrated that BHRF1 disturbs mitochondrial dynamics and subsequently stimulates mitophagy, a cellular process that can specifically sequester and degrade mitochondria by autophagy, leading to the inhibition of type I IFN response by EBV (Vilmen *et al*., 2020). Here, we explored the role of the MT network in the innate immunity escape induced by BHRF1 and we revealed the contribution of the MT hyperacetylation on the BHRF1-induced mitochondrial clustering and on the BHRF1-mediated counteraction of the IFN responses.

## Results

### BHRF1-induced mito-aggresome is organized around the centrosome

The EBV-encoded BHRF1 localizes to mitochondria, and its expression in HeLa cells dramatically alters mitochondrial dynamics and distribution, (Vilmen *et al*., 2020). By confocal microscopy, we observed abnormal mitochondrial clusters close to the nucleus in BHRF1-expressing cells, whereas in control cells (empty vector [EV] transfection), mitochondria remained distributed throughout the cytoplasm (***Figure 1A***). This mitochondrial phenotype, known as “mito-aggresome”, was in accordance with our previous observations. To quantify this phenotype, we arbitrarily considered that, when the compaction index (CI) of mitochondria is above 0.4, the cell presents a mito-aggresome, whereas when the CI value is below 0.4, mitochondria are distributed homogeneously throughout the cell cytoplasm (Narendra *et al*., 2010; Vilmen *et al*., 2020). The expression of BHRF1 significantly increased the CI values, which corresponds to the formation of a mito-aggresome in almost 80% of the cells (***Figure 1B***). To confirm the role of BHRF1 in the context of EBV infection, we analyzed the mitochondrial phenotype in EBV-positive Akata B cells, during latency or after treatment with anti-human IgG to induce EBV reactivation (***Figure 1 – figure supplement 1***). We clearly observed mito-aggresomes upon EBV reactivation, whereas the knockdown (KD) of BHRF1 completely blocked their formation. Since BHRF1 is associated with mitochondrial fragmentation (***Figure 1A - insets***), we precisely evaluated mitochondrial shape by calculation of the aspect ratio (AR) and the form factor (FF), two parameters that reflect the mitochondrial length and the branching of mitochondria, respectively. As calculated in ***Figure 1C***, BHRF1 expression significantly reduced both AR and FF parameters, confirming the induction of mitochondrial fission.

**Figure 1.**
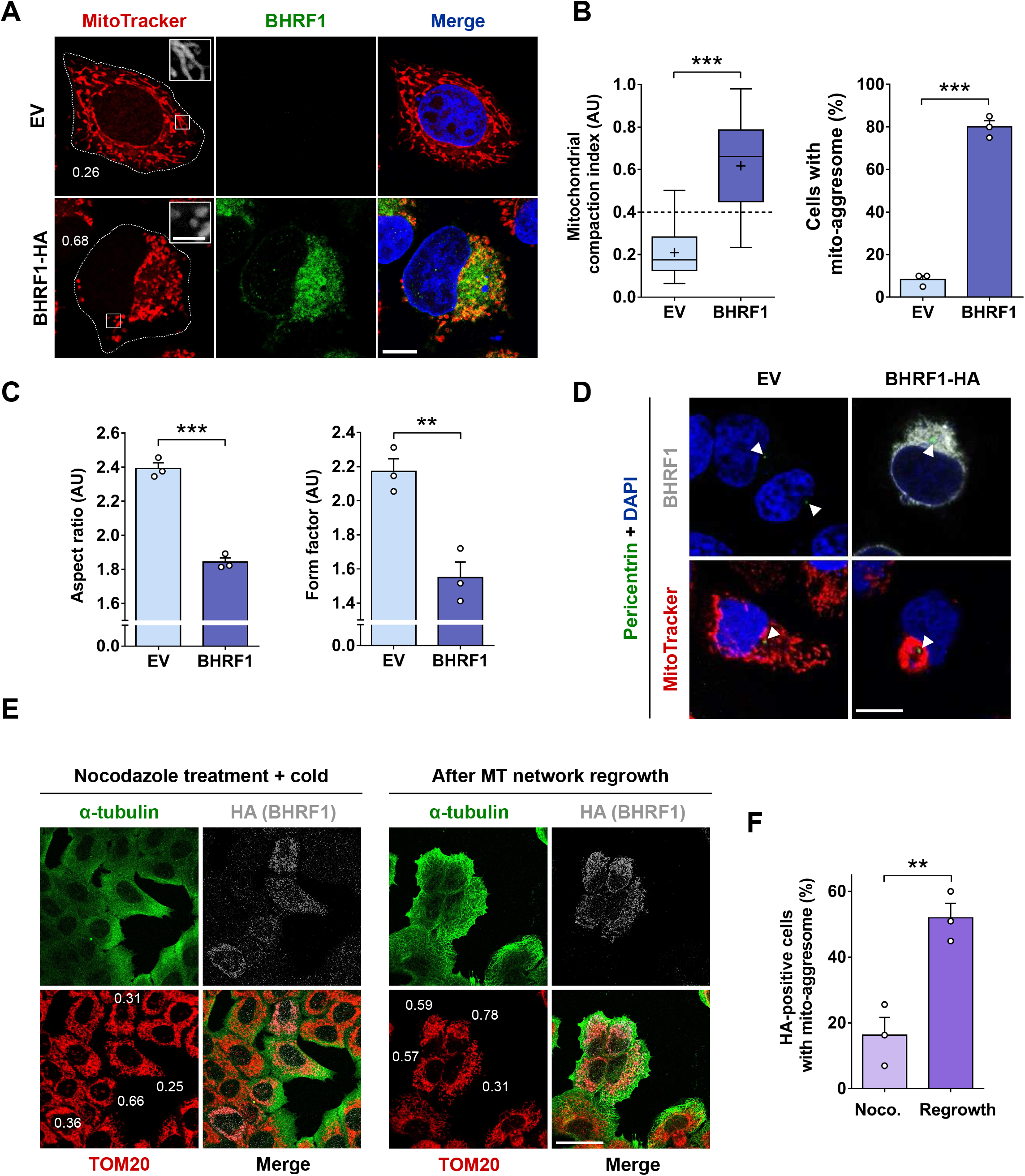
BHRF1 induces the formation of mito-aggresomes in a MT-dependent manner. (**A-C**) HeLa cells were transfected for 24 h with BHRF1-HA plasmid or with EV as a control. Mitochondria were labeled with MitoTracker and cells immunostained for BHRF1. Nuclei were stained with DAPI. (**A**) Confocal images with insets (3X) on mitochondrial phenotype. Values of mitochondrial CI are indicated on representative cells. Scale bars: 10 μm and 2 μm for insets. (**B**) *Left*, assessment of mitochondrial aggregation by calculation of CI. *Right*, percentage of cells presenting a mito-aggresome (n = 20 cells per condition). (**C**) Quantification of mitochondrial fission parameters, aspect ratio (AR; *left panel*) and form factor (FF, *right panel*). (**D**) Confocal images of HeLa cells transfected with BHRF1-HA plasmid (or EV) and immunostained for pericentrin and BHRF1 (*upper panel*) or mitochondria (*lower panel*). Nuclei were stained with DAPI. Scale bar: 20 μm. Arrowheads show the centrosomal localization of pericentrin. (**E-F**) MT network depolymerization assay. (**E**) Representative images where BHRF1 expression was visualized by HA immunostaining (gray), the MT network by α-tubulin immunostaining (green), and mitochondria are immunostained with an anti-TOM20 antibody (red). Values of mitochondrial CI are indicated on representative cells. Scale bars: 10 μm. (**F**) Percentage of BHRF1-HA-positive cells presenting a mito-aggresome (n = 30 cells per condition) in cells treated with nocodazole and after MT regrowth. Data represent the mean ± SEM of three independent experiments. ** P < 0.01; *** P < 0.001 (Student’s t-test).

As mitochondria localize near the nucleus upon BHRF1 expression, the involvement of the MT network in this mitochondrial redistribution was investigated. In mammalian cells, mitochondria move along cytoskeletal tracks to sites of high-energy demand in an MT-dependent manner (Detmer and Chan, 2007). At interphase, the MT network radiates from the centrosome, the main MT organizing center for directing the polarity and the orientation of MTs. By co-staining BHRF1-expressing cells for mitochondria and pericentrin, a classical centrosomal marker, we observed that mitochondria are concentrated around the centrosome to form a mito-aggresome (***Figure 1D***). Moreover, a 3D reconstruction confirmed the clustering of BHRF1 in the vicinity of the nucleus around the centrosome (***Figure 1 – figure supplement 2***).

### The MT network is required for BHRF1-induced mito-aggresome formation

To test whether MTs play a role in the mitochondrial redistribution induced by BHRF1 expression, the whole MT network was disassembled using a treatment mixing nocodazole, a classical MT-destabilizing drug, and exposure to cold (Geeraert *et al*., 2010). As shown in ***Figure 1E - left panel*** this treatment disassembled MTs (fuzzy staining of α-tubulin). The removal of nocodazole followed by 1 hour in a warm complete medium at 37°C restored the MT network (***Figure 1E - right panel***). In cells without BHRF1 expression, mitochondria were homogeneously distributed in the cytoplasm, and the disorganization of the MT network had no impact on their distribution. Conversely, when MTs were depolymerized, the mito-aggresome formation was abolished (***Figure 1E - left panel***) and mitochondria were then distributed throughout the cell (CI values mainly below 0.4). This phenomenon was reversible since the mitochondria rapidly re-aggregated to different extents within 1 hour after reassembly of the MT network (most of the CI values above 0.4) (***Figure 1E - right panel***). In contrast, mitochondria in non-transfected neighboring cells remained scattered within the cytoplasm after nocodazole removal. The quantification of mito-aggresomes in BHRF1-positive cells showed that whereas less than 20% of the cells presented mitochondrial clustering after depolymerization treatment (***Figure 1F***), this proportion went back to more than 50% after repolymerization of the MTs (compare to 80% without treatment). We concluded that the MT network is required for the mito-aggresome formation.

The actin network is also known to be involved in mitochondrial transport and fission (De Vos *et al*.,2005; López-Doménech *et al*., 2018). We thus checked if F-actin disorganization affects the mitochondrial phenotype induced by BHRF1. To do so, we treated HeLa cells with cytochalasin B to disrupt F-actin but we did not observe any changes in the mitochondrial phenotype in BHRF1-positive cells (***Figure 1 – figure supplement 3A***). BHRF1 still induced mito-aggresome formation and fission at the same level as in the control (***Figure 1 – figure supplement 3B***). Therefore, the actin network does not seem to be involved in the mitochondrial phenotype.

### EBV reactivation induces BHRF1-dependent MT hyperacetylation

We next explored whether BHRF1 could modify the MT network by assessing α-tubulin acetylation level. Interestingly, upon BHRF1 expression, whereas α-tubulin staining was unchanged, acetyl-α-tubulin was redistributed and colocalized with BHRF1 (***Figure 2A***). Moreover, we observed a brighter signal of acetyl-α-tubulin in BHRF1 expressing cells, suggesting an increase in α-tubulin acetylation. This was confirmed by western-blot analysis showing an increase of 60% under BHRF1 expression (***Figure 2B***).

**Figure 2.**
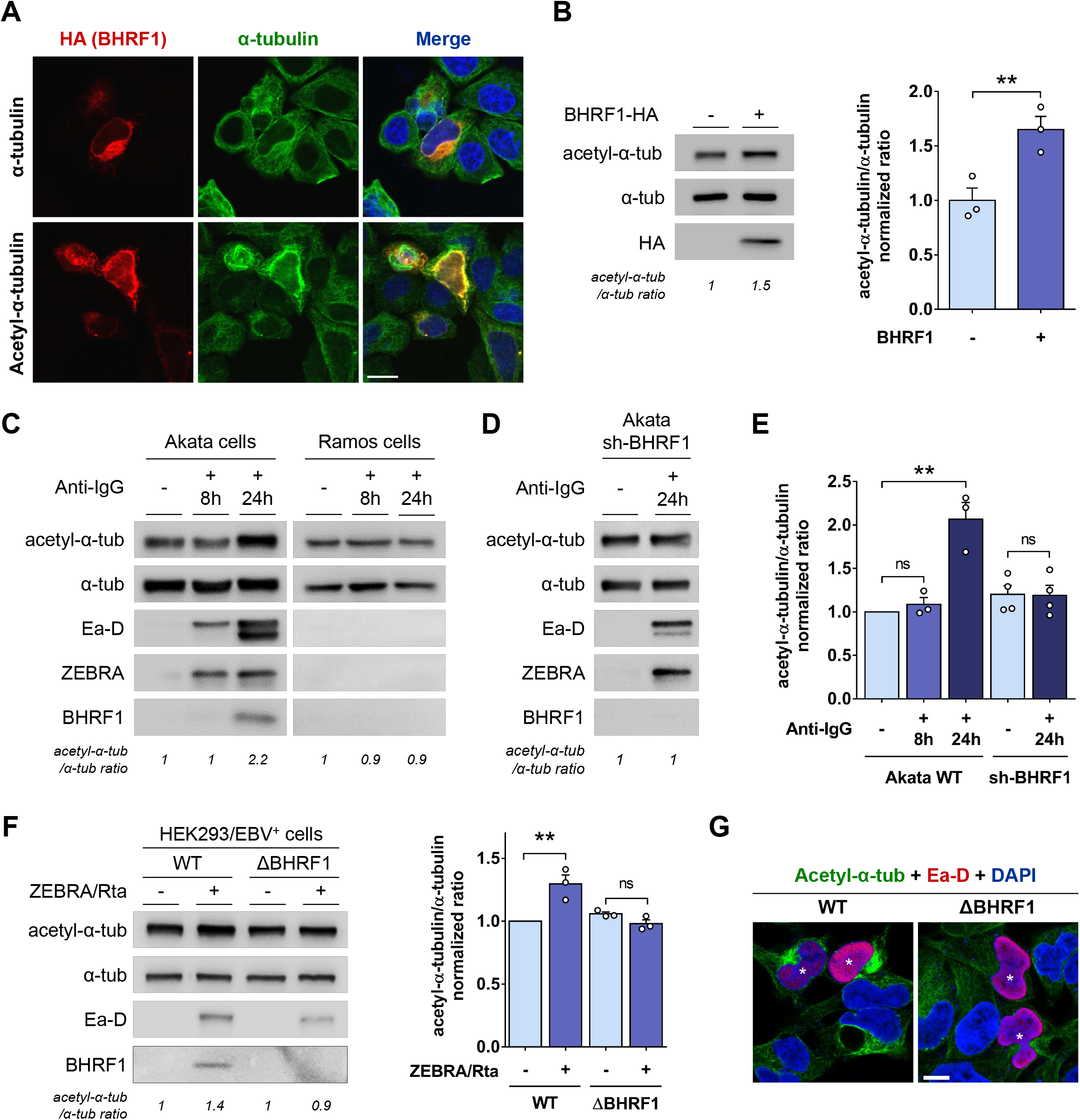
BHRF1 stimulates MT hyperacetylation. (**A**) Representative images of HeLa cells expressing BHRF1-HA and immunostained for HA and α-tubulin (*upper panel*) or acetyl-α-tubulin (*lower panel*). Nuclei were stained with DAPI. Scale bar: 20 μm. (**B**) *Left*, immunoblot analysis of acetyl and total α-tubulin in BHRF1-HA-transfected HeLa cells. *Right*, normalized ratios of acetyl-α-tubulin to total α-tubulin. (**C**) Immunoblot analysis of acetyl-α-tubulin, α-tubulin, Ea-D, ZEBRA and BHRF1 in the Akata and Ramos cells treated or not with anti-human IgG for 8 h or 24 h to induce EBV reactivation. (**D**) Immunoblot analysis of acetyl-α-tubulin, α-tubulin, Ea-D, ZEBRA and BHRF1 in Akata cells deficient for BHRF1 expression (sh-BHRF1). (**E**) Normalized ratios of acetyl-α-tubulin to α-tubulin in Akata cells. (**F-G**) EBV WT or EBV ΔBHRF1 were reactivated in HEK293 cells by transfection with ZEBRA and Rta plasmids for 24 h. (**F**) *Left*, immunoblot analysis of acetyl-α-tubulin, α-tubulin, Ea-D and BHRF1. *Right*, normalized ratios of acetyl-α-tubulin to α-tubulin. (**G**) Confocal images of HEK293/EBV+ cells immunostained for acetyl-α-tubulin and Ea-D. Nuclei were stained with DAPI. Scale bar: 20 μm. Stars indicate EBV-reactivated cells. Data represent the mean ± SEM of three independent experiments. ns = non-significant; ** P < 0.01 (Student’s t-test).

To confirm these results in the context of EBV infection, we analyzed the level of MT acetylation in EBV-positive Akata B cells (***Figures 2C and 2E***). We used, as a control, EBV-negative Ramos B cells similarly treated with anti-IgG. We observed an increase of MT acetylation only 24h after EBV reactivation in Akata cells (visualized by Ea-D and ZEBRA viral protein expression). This MT hyperacetylation coincided with BHRF1 expression. We confirmed that BHRF1 is required for this phenotype in the context of EBV infection of B cells, using an shRNA targeting BHRF1 (***Figures 2D and 2E***). In parallel, in HEK293/EBV+ epithelial cells, the EBV genomes WT and ΔBHRF1 were reactivated by co-transfection of plasmids encoding the trans-activator proteins ZEBRA and Rta (***Figures 2F and 2G***). We similarly observed a BHRF1-dependent MT hyperacetylation upon EBV reactivation in epithelial cells. Due to a low EBV reactivation rate in this model, the increase of MT hyperacetylation observed, although statistically significant, was less important than in Akata cells. However, by confocal microscopy, the increase of tubulin acetylation was clearly seen in WT HEK293/EBV+ and not in ΔBHRF1 (***Figure 2G***).

### MT hyperacetylation is required for the mito-aggresome formation

To determine whether this hyperacetylation could be involved in BHRF1-induced mitochondrial alterations, we expressed in HeLa cells a non-acetylatable α-tubulin mutant (mCherry-α-tubulin K40A) which prevents MT hyperacetylation (Dompierre *et al*., 2007). Expression of this non-acetylatable α-tubulin mutant does not alter the MT network architecture and this protein is incorporated in the whole MT network (Geeraert *et al*., 2010). However, the whole MT network hyperacetylation is prevented, since α-tubulin acetylation only occurs on polymers and the MT network is totally decorated with this non-acetylatable α-tubulin. We thus verified by immunoblot analysis that BHRF1 was unable to induce hyperacetylation of α-tubulin when the α-tubulin K40A mutant was expressed (***Figure 3 – figure supplement 1A***). Then, the impact of α-tubulin K40A on the clustering of mitochondria was analyzed by confocal microscopy (***Figures 3A and 3B***). In cells co-expressing α-tubulin K40A and BHRF1, CI mean value was below 0.4 and mitochondria were homogeneously distributed throughout the cytoplasm, similarly to control cells. Thus, we concluded that BHRF1-induced mito-aggresome formation depends on MT hyperacetylation. This was confirmed in the context of EBV reactivation (***Figures 3C and 3D***). EBV reactivation was visualized in HEK293/EBV WT cells using the staining of Ea-D viral protein.

**Figure 3.**
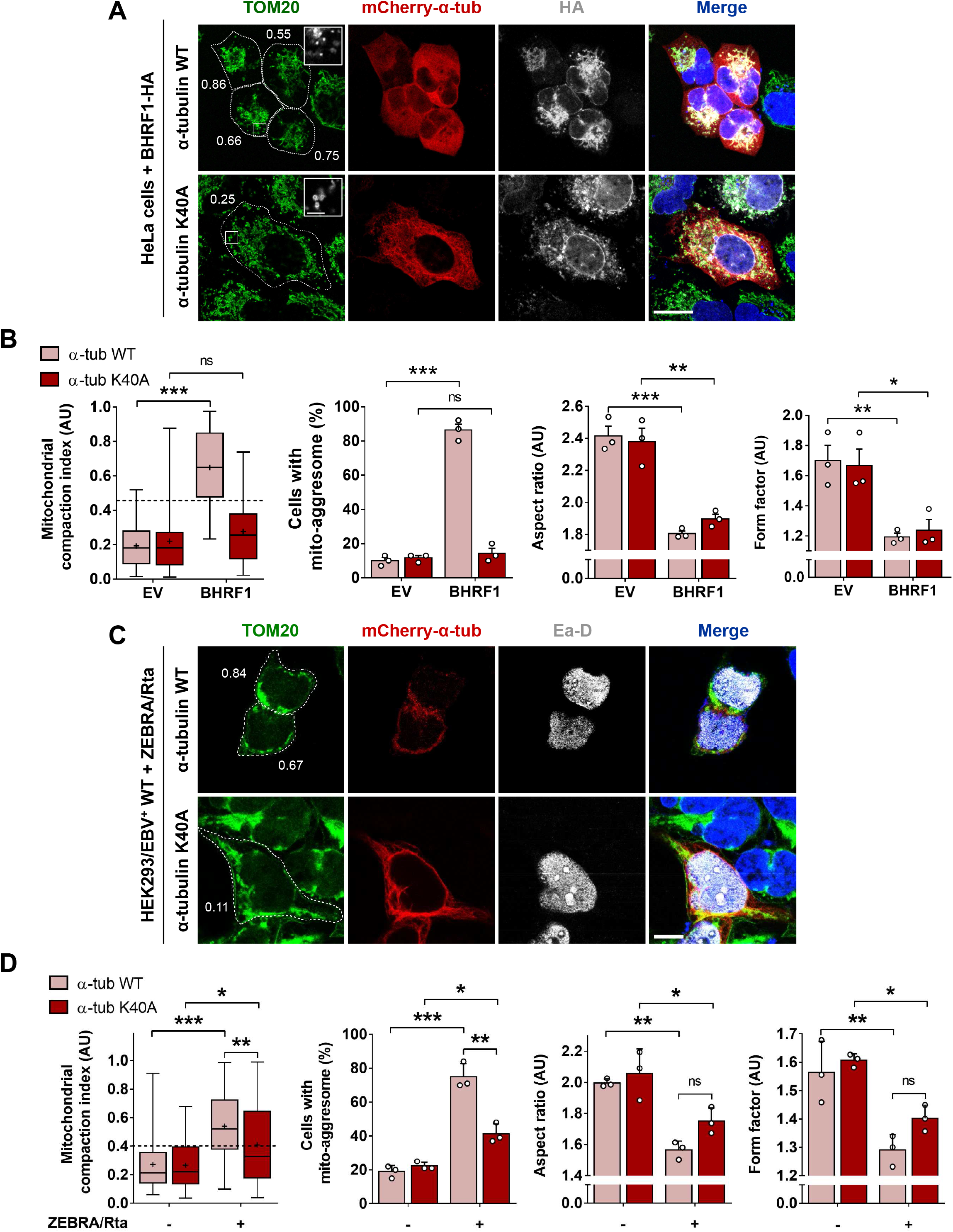
MT hyperacetylation is required for BHRF1-induced mito-aggresomes formation. (**A-B**) HeLa cells were co-transfected with plasmids encoding BHRF1-HA and either mCherry-α-tubulin WT or a non-acetylatable form (K40A). (**A**) Confocal images with insets (3X) of cells immunostained for HA and TOM20. Nuclei were stained with DAPI. Scale bars: 10 μm and 4 μm for insets. Values of mitochondrial CI are indicated on representative cells. Images of control cells are presented in **Figure 3 – figure supplement 1B**. (**B**) Quantification of CI, percentage of cells with a mito-aggresome and mitochondrial fission parameters (n = 20 cells per condition). (**C-D**) HEK293/EBV^+^ WT cells were co-transfected with plasmids encoding ZEBRA, Rta and either mCherry-α-tubulin WT or K40A. (**C**) Confocal images of cells immunostained for Ea-D and TOM20. Nuclei were stained with DAPI. Scale bar: 10 μm. Values of mitochondrial CI are indicated on representative cells. (**D**) Quantification of CI, percentage of cells with a mito-aggresome and mitochondrial fission parameters (n = 20 cells per condition). Data represent the mean ± SEM of three independent experiments. ns = non-significant; * P < 0.05; ** P < 0.01; *** P < 0.001 (Student’s t-test).

Interestingly, we observed that without α-tubulin hyperacetylation, BHRF1 expression or EBV reactivation were still able to promote mitochondrial fragmentation (***Figures 3B and 3D***). Confocal images (***Figure 3A, insets***) clearly showed that mitochondria were drastically smaller upon BHRF1 expression in both WT and K40A conditions. We previously reported that knockdown of Drp1 (dynamin-related protein 1), a master regulator of mitochondrial fission, prevents the BHRF1-induced mitochondrial phenotype (Vilmen *et al*., 2020). We thus investigated the role of Drp1 in MT hyperacetylation and found that the loss of Drp1 inhibits MT hyperacetylation induced by BHRF1 expression (***Figure 3 – figure supplement 2A***). In the same way, treatment of Akata cells with Mdivi-1, which inhibits mitochondrial fission, blocked MT hyperacetylation induced by EBV reactivation (***Figure 3 – figure supplement 2B***). Taken together, these results demonstrated that the hyperacetylation of MTs requires mitochondrial fission and is essential for BHRF1-induced mito-aggresome formation.

### MT hyperacetylation is necessary for the inhibition of type I IFN response by BHRF1

Mitochondria appear to be central organelles for the induction of antiviral innate immunity. Our group and others previously demonstrated that impairing the mitochondrial morphodynamics leads to the inhibition of IFN response (Castanier *et al*., 2010; Chatel-Chaix *et al*., 2016; Vilmen *et al*., 2020; Yoshizumi *et al*., 2014). Considering the importance of tubulin hyperacetylation for the formation of mito-aggresomes, we investigated whether MT hyperacetylation was required for the inhibition of type I IFN response by BHRF1. We first performed an IFN-β promoter-driven luciferase reporter assay in BHRF1-expressing HEK293T cells (***Figure 4A***). The activation of the IFN-signaling pathway was induced by transfection of an active form of RIG-I, designated ΔRIG-I (2xCARD) (Yoneyama *et al*., 2004), which constitutively activates the MAVS-dependent pathway through CARD domains interaction. As previously observed, BHRF1 dramatically blocked the IFN-β promoter activation. By contrast, the expression of the α-tubulin K40A significantly prevented this inhibition (***Figure 4A***). To confirm our results, we next examined the cellular localization of IRF3 by immunofluorescence. In basal conditions, IRF3 staining is diffuse throughout the cytoplasm. When ΔRIG-I (2xCARD) is expressed, IRF3 translocates from the cytoplasm into the nucleus but the expression of BHRF1 blocked this translocation (***Figure 4B***). However, when MT hyperacetylation was prevented, BHRF1 was not able to block IRF3 nuclear translocation (***Figure 4B***). Quantification of this assay showed that expression of the non-acetylatable form of α-tubulin totally restored IRF3 nuclear translocation upon BHRF1 expression (***Figure 4C***). Therefore, our results demonstrated that MT hyperacetylation is required for BHRF1 to inhibit type I IFN response.

**Figure 4.**
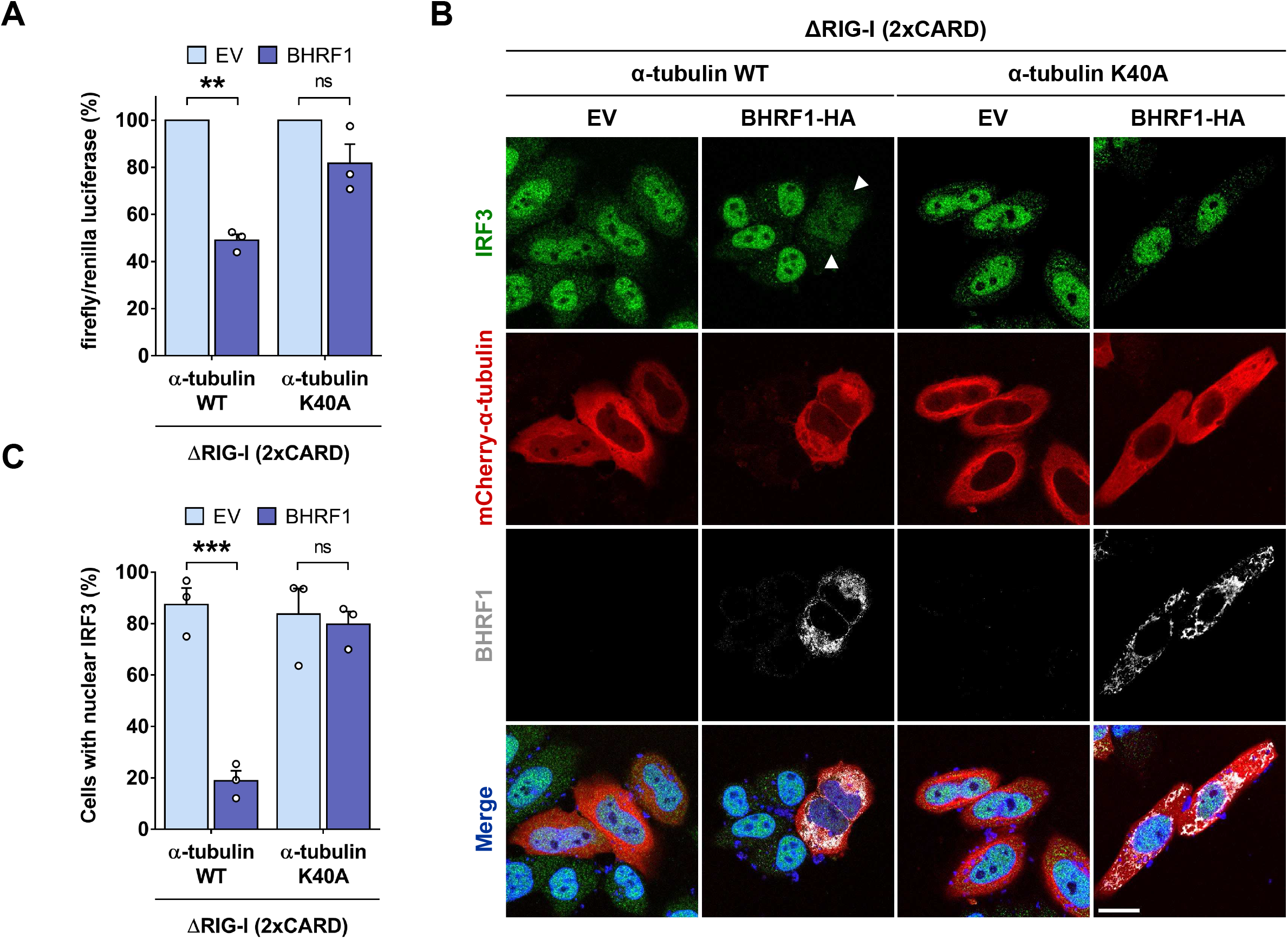
MT hyperacetylation is required for BHRF1 to block IFN. (**A**) Luciferase reporter assay on HEK293T cells co-expressing BHRF1 (or EV) and α-tubulin K40A (or WT). Activation of the IFN-β promoter was analyzed 24 h post-transfection. Firefly/renilla luciferase ratios were calculated and normalized to control conditions. (**B-C**) HeLa cells were co-transfected for 24 h with plasmids encoding BHRF1-HA (or EV), mCherry-α-tubulin K40A (or WT), and ΔRIG-I (2xCARD). (**B**) Confocal images. BHRF1-HA-transfected cells were visualized with an anti-HA antibody and cells were immunostained with an-anti IRF3 antibody. Nuclei were stained with DAPI. Scale bar: 20 μm. Arrows indicate BHRF1-expressing cells without IRF3 nuclear localization. (**C**) Percentage of cells expressing mCherry-α-tubulin and presenting IRF3 nuclear localization (n = 50 cells per condition). Data represent the mean ± SEM of three independent experiments. ns = non-significant; ** P < 0.01; *** P < 0.001 (Student’s t-test).

We investigated whether this strategy of IFN inhibition through MT hyperacetylation was original and specific to BHRF1. To explore this, we assayed the impact of three different mitophagy inducers, AMBRA1-ActA, Oligomycin/Antimycin A (O/A) and carbonyl cyanide m-chlorophenyl hydrazone (CCCP), on the interplay between tubulin acetylation and IFN inhibition (Lazarou *et al*., 2015; Narendra *et al*., 2008; Strappazzon *et al*., 2015) (***Figure 4 – figure supplement 1***). Similar to BHRF1, expression of AMBRA1-ActA induced mitochondrial fission and mito-aggresome formation, leading to IFN inhibition (***Figure 4 – figure supplement 1A and 1E***). However, AMBRA1-ActA did not impact tubulin acetylation (***Figure 4 – figure supplement 1B***). On the contrary, the mitophagy inducers O/A and CCCP did not induce mito-aggresome formation, although CCCP increased mitochondrial fission (***Figure 4 – figure supplement 1C***). These two treatments were able to reduce IFN production, through mitophagy induction. Indeed, treatment with chloroquine (CQ), which blocks autophagic degradation, restored the IFN level (***Figure 4 – figure supplement 1E***). Similar to AMBRA1-ActA, neither O/A nor CCCP impacted tubulin acetylation (***Figure 4 – figure supplement 1D***). These data, summarized in ***Figure 4 – figure supplement 1F***, clearly showed that these three mitophagy inducers blocks IFN response, but without regulating MT acetylation level. This demonstrates the originality of BHRF1 strategy to inhibit IFN but also the fact that more generally the induction of mitophagy can impede IFN production.

### BHRF1 requires acetyltransferase ATAT1 to modify MTs and form mito-aggresomes

Thereafter, we wanted to characterize the mechanism of MT hyperacetylation induction by BHRF1. Tubulin acetylation results from the balance between the activities of tubulin acetyltransferases and deacetylases. Therefore, a hyperacetylation of the MTs can be the consequence of either the inhibition of deacetylases or the activation of an acetyltransferase. First of all, we explored a potential inhibitory role of BHRF1 on the two main α-tubulin deacetylases, HDAC6 (histone deacetylase 6) and SIRT2 (sirtuin type 2) (Hubbert *et al*., 2002; North *et al*., 2003). Indeed, it has been observed that KD of SIRT2 results in an aberrant mitochondrial distribution in the perinuclear region (Misawa *et al*., 2013), and another study showed that the depletion of SIRT2 results in nuclear envelope shape defects (Kaufmann *et al*., 2016). These observations are clearly similar to the effect of BHRF1 on mitochondria, as well as its distribution around the nuclear envelope and the modification of the nucleus shape (***Figure 1A***). We thus performed an *in vitro* tubulin deacetylation assay (Mackeh *et al*., 2014). This test consists of mixing the highly acetylated porcine brain tubulin with lysates of BHRF1-expressing cells. Lysate of control EV-transfected cells reduced the acetylation of tubulin by approximately 50% compared to the condition without cell lysate (***Figure 5A***). The same deacetylase activity was observed with BHRF1-positive lysates, indicating that BHRF1 does not inhibit the deacetylases.

**Figure 5.**
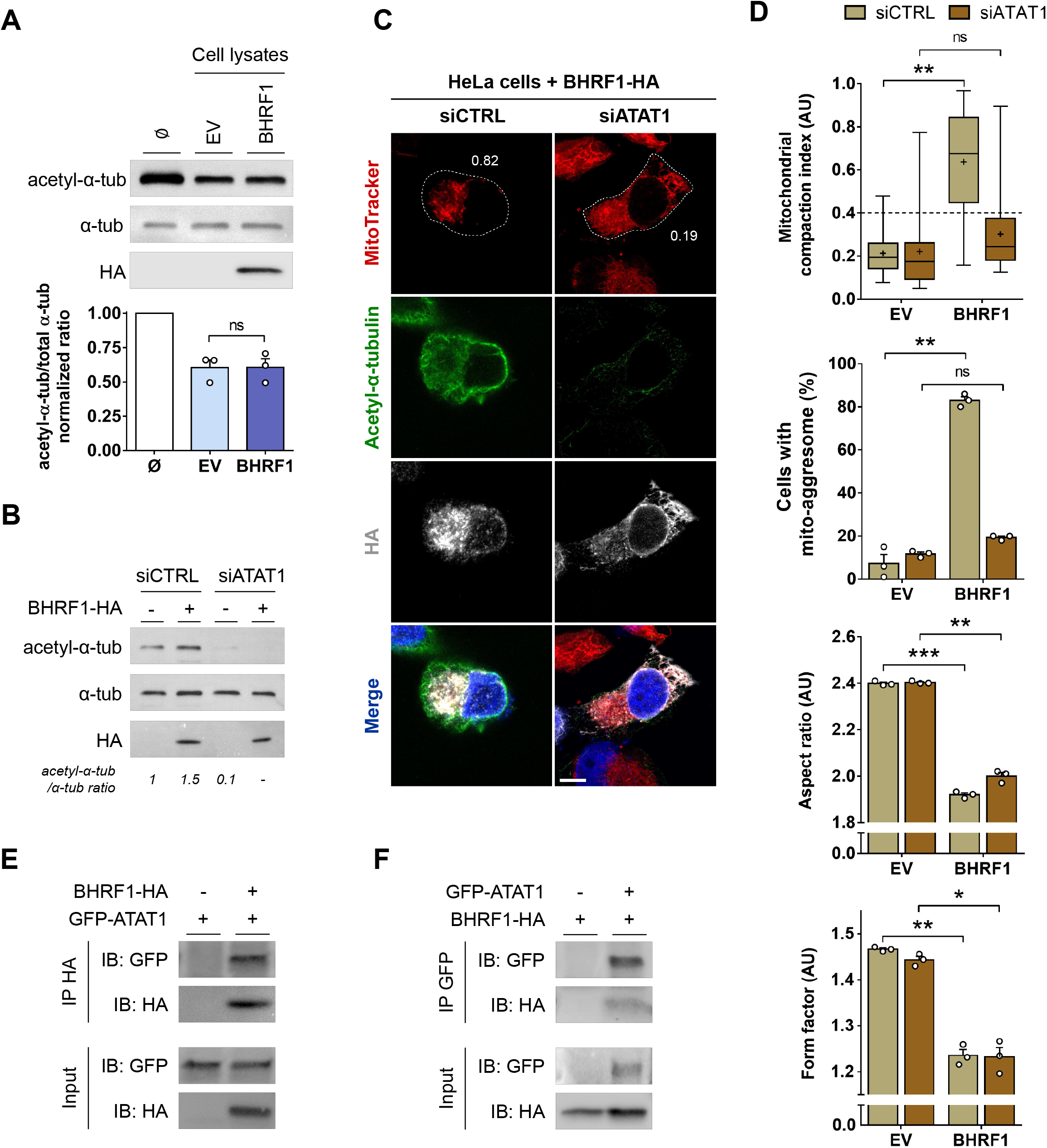
MT hyperacetylation and mito-aggresome formation require ATAT1, which interacts with BHRF1. (**A**) Tubulin deacetylation assay. HeLa cells were transfected with BHRF1-HA plasmid (or EV) and lysed 48 h post-transfection. Cell lysates were incubated in vitro with porcine brain tubulin and NAD+. BHRF1-HA, acetyl-α-tubulin and α-tubulin were detected by immunoblot, and the histograms represent the normalized ratios of acetyl-α-tubulin to α-tubulin. (**B-D**) After knockdown of ATAT1 by siRNA transfection, HeLa cells were transfected with BHRF1-HA for 24 h. (**B**) Loss of ATAT1 activity was visualized by immunoblot analysis of acetyl-α-tubulin and α-tubulin. (**C**) Confocal images. Mitochondria were labeled with MitoTracker, and cells were immunostained for acetyl-α-tubulin and HA. Nuclei were stained with DAPI. Values of mitochondrial CI are indicated on representative cells. Scale bar: 20 μm. Images of EV-transfected cells are presented in ***Figure 5 – figure supplement 1***. (**D**) Quantification of CI, percentage of cells with a mito-aggresome and mitochondrial fission parameters (n = 20 cells per condition). (**E-F**) HeLa cells were co-transfected with GFP-ATAT1 and BHRF1-HA plasmids for 24 h. (**E**) After immunoprecipitation of BHRF1 with an anti-HA antibody, proteins were detected by immunoblotting with anti-GFP and anti-HA antibodies. (**F**) After immunoprecipitation of GFP-ATAT1 with an anti-GFP antibody, proteins were detected by immunoblotting with anti-GFP and anti-HA antibodies. Data represent the mean ± SEM of three independent experiments. ns = non-significant; * P < 0.05; ** P < 0.01; *** P < 0.001 (Student’s t-test).

We subsequently investigated the role of ATAT1 (alpha-tubulin N-acetyltransferase 1) on BHRF1-induced MT hyperacetylation (Shida *et al*., 2010). We knocked down ATAT1 expression by a siRNA approach and observed no more detectable acetylation activity under BHRF1 expression (***Figure 5B***), suggesting the involvement of ATAT1 in BHRF1-induced hyperacetylation. Then, we checked the mitochondrial network and observed that ATAT1 KD prevents mito-aggresome formation but not mitochondrial fission (***Figures 5C and 5D***). These results led us to conclude that BHRF1 activates the acetyltransferase ATAT1, leading to MT hyperacetylation and subsequent mito-aggresome formation. To go further, we analyzed the distribution of GFP-tagged ATAT1 in presence of BHRF1 in HeLa cells (***Figure 5 – figure supplement 2***) and we observed a clear colocalization between GFP-ATAT1 and BHRF1 and, in all cells expressing GFP-ATAT1, a predictable increase in tubulin acetylation. Finally, co-immunoprecipitation assays revealed that BHRF1 interacts with ATAT1 in HeLa cells (***Figures 5E and 5F***). Our results suggest that the interaction between BHRF1 and ATAT1 leads to MT hyperacetylation.

### Mito-aggresome formation requires retrograde transport of mitochondria via dyneins

We observed in ***Figure 1E*** that after removal of nocodazole and regrowth of the MT network, mitochondria rapidly re-aggregated in BHRF1-expressing cells, suggesting that an MT-dependent motor activity is involved in the mito-aggresome formation. The hyperacetylation of MTs has been previously shown to facilitate the recruitment and binding of MAPs on the tubulin (Dompierre *et al*., 2007). We therefore hypothesized that BHRF1-induced mitochondrial clustering next to the nucleus could use dyneins. To investigate this hypothesis, we inhibited dynein functions by overexpression of p50-dynamitin. The overexpression of one subunit of the dynactin complex (p50-dynamitin) disrupts it and results in the dissociation of dyneins from the MT network. We observed that the expression of this construction totally prevents mito-aggresome formation in BHRF1-expressing cells (***Figures 6A and 6B***). Moreover, the ability of BHRF1 to induce mitochondrial fission was decreased when dyneins were inhibited, as we observed a partial but significant restoration of mitochondrial fission parameters (AR and FF; ***Figure 6B***).

**Figure 6.**
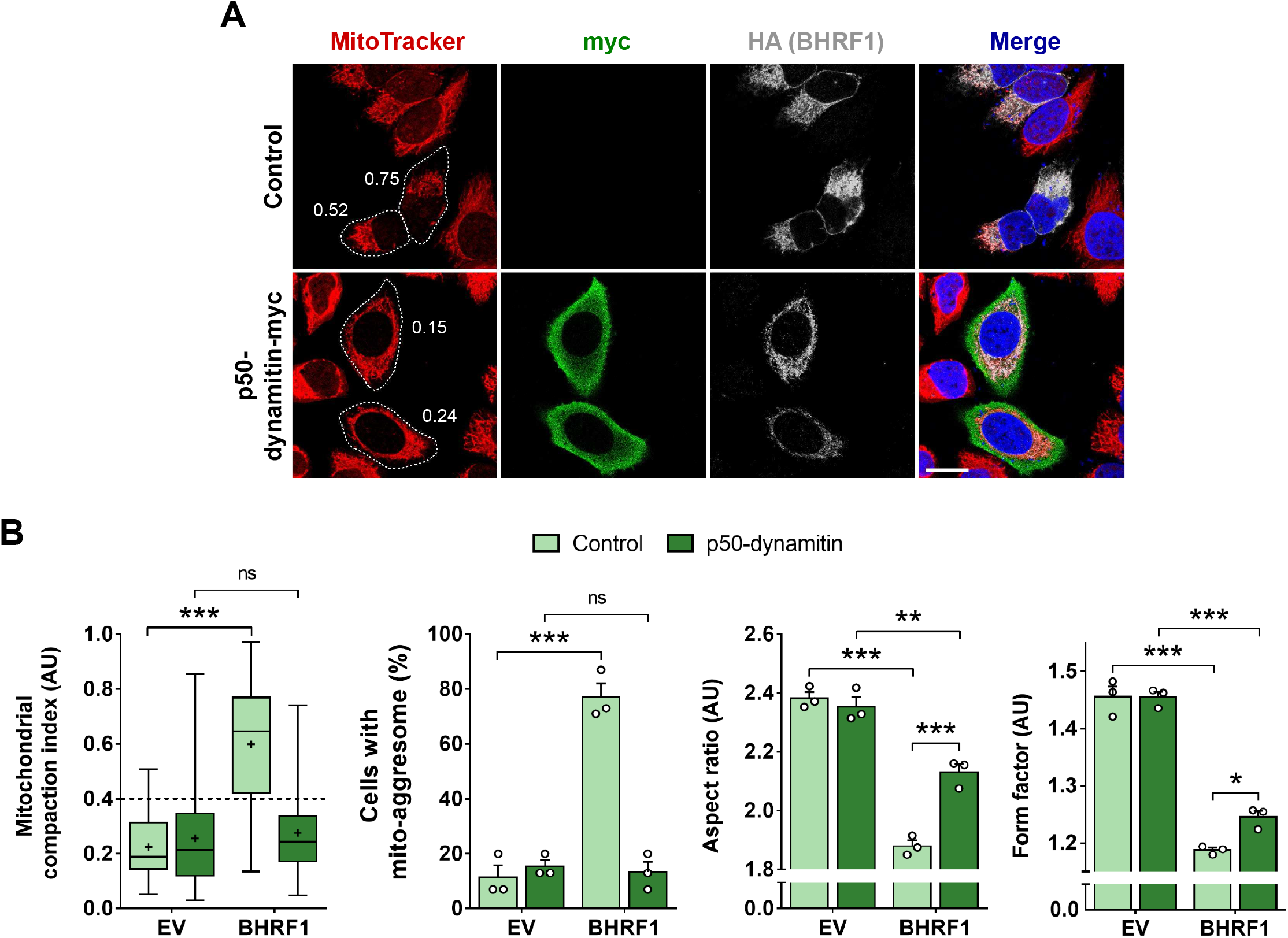
Dynein-based transport is required for BHRF1 to aggregate mitochondria next to the nucleus. (**A-B**) HeLa cells were co-transfected with plasmids encoding BHRF1-HA and p50-dynamitin-myc (or control) for 24 h. (**A**) Confocal images. Mitochondria were labeled with MitoTracker, and cells were immunostained for c-myc and HA. Nuclei were stained with DAPI. Values of mitochondrial CI are indicated on representative cells. Scale bar: 20 μm. Images of cells co-expressing EV and p50-dynamitin-myc are presented in **Figure 6 – figure supplement 1A**. (**B**) Quantification of CI, percentage of cells with a mito-aggresome and mitochondrial fission parameters (n = 20 cells per condition). Data represent the mean ± SEM of three independent experiments. ns = non-significant; * P < 0.05, ** P < 0.01; *** P < 0.001 (Student’s t-test).

To confirm the importance of dynein motors in the mito-aggresome formation, we treated cells with ciliobrevin-D, a specific inhibitor of dyneins, which blocks their gliding along MTs and their ATPase activity. As with overexpression of p50-dynamitin, BHRF1 was not able to induce mito-aggresomes in cells treated with ciliobrevin-D (***Figure 6 – figure supplement 1B and 1C***). However, BHRF1 still induced a slight mitochondrial fission phenomenon. Altogether, these results demonstrated that fragmented mitochondria are clustered next to the nucleus in a dynein-dependent manner.

### MT hyperacetylation does not affect BHRF1-pro-autophagic activity, but is required for the first step of mitophagy induction

Our group previously demonstrated that BHRF1 first stimulates autophagy and then induces mitophagy, the selective degradation of mitochondria (Vilmen *et al*., 2020). Autophagy is a conserved cellular process allowing the removal and recycling of damaged or supernumerary cellular components. This process involves double-membrane vesicles, called autophagosomes, which fuse with lysosomes to degrade the content. Selective autophagy consists of the recognition of a specific cargo by a molecular receptor that links the cargo to the autophagosome for its removal (Green and Levine, 2014).

Several studies demonstrated the role of MTs during the autophagic process, notably during the formation of the autophagosomes (Geeraert *et al*., 2010; Köchl *et al*., 2006; Mackeh *et al*., 2013). Thus, we investigated whether BHRF1 required MT hyperacetylation to induce autophagy. The autophagic flux was studied in HeLa cells co-expressing α-tubulin K40A and BHRF1. We first analyzed by immunofluorescence the accumulation of LC3 (microtubule-associated protein 1 light chain 3), a classical marker of autophagosomes in the presence or absence of CQ, an inhibitor of the autophagic flux. Expression of BHRF1 induced a clear accumulation of LC3 dots in the cytoplasm, and even more after CQ treatment, when non-acetylatable α-tubulin was expressed (***Figure 7A***). We quantified LC3 dots and we observed no difference when MTs were hyperacetylated or not (***Figure 7B and Figure 7 – figure supplement 1B***). We confirmed the result by analyzing the accumulation of lipidated LC3 (LC3-II) by immunoblot and we observed an increase in LC3-II levels in BHRF1-expressing cells, independently of MT hyperacetylation (***Figure 7C and Figure 7 – figure supplement 1C***). This increase is potentiated after CQ treatment. Therefore, we concluded that BHRF1 stimulates the biogenesis of autophagosomes and the autophagic flux independently of MT hyperacetylation.

**Figure 7.**
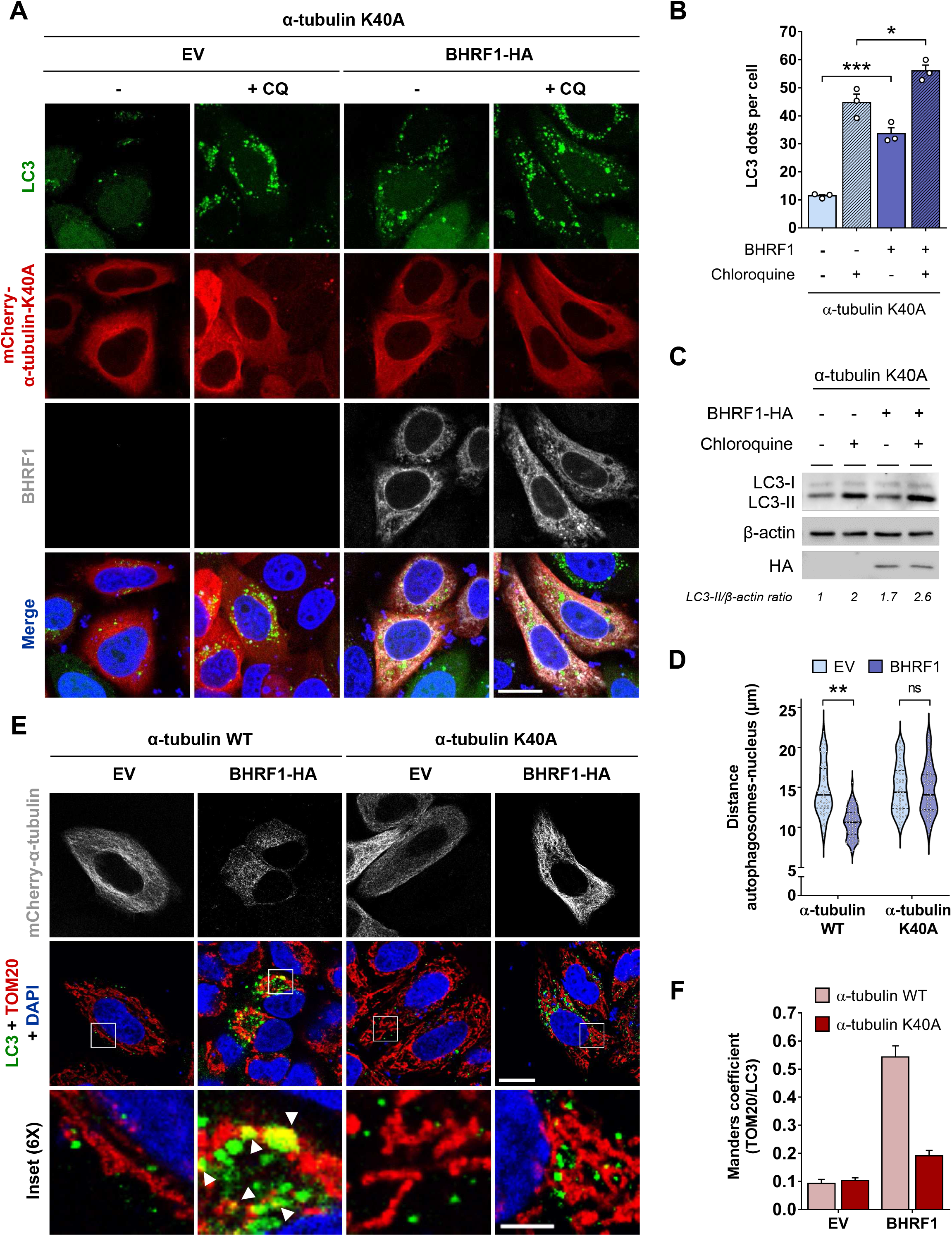
MT hyperacetylation is dispensable for BHRF1-pro-autophagic activity, but essential for mitophagy. HeLa cells were co-transfected for 24 h with plasmids encoding BHRF1-HA (or EV) and mCherry-α-tubulin K40A (or WT) and treated with chloroquine when indicated. (**A**) Confocal images. Cells were immunostained for BHRF1 and LC3 and nuclei stained with DAPI. Scale bar: 10 μm. Images of cells transfected with mCherry-α-tubulin WT are presented in **Figure 7 – figure supplement 1A**. (**B**) Quantification of LC3 dots (n = 30 cells per condition). (**C**) Immunoblot analysis of LC3 and BHRF1-HA expression. β-actin was used as a loading control. (**D**) Measurement of the distance between autophagosomes and nucleus (n = 60 cells from three independent experiments). (**E**) Confocal images of cells immunostained for TOM20 and LC3. Nuclei were stained with DAPI. Insets (6X) show colocalization events (see arrows). Scale bars: 10 μm and 5 μm for insets. (**F**) Colocalization level (Manders coefficient) between mitochondria (TOM20) and autophagosomes (LC3) (n = 30 cells per condition). Data represent the mean ± SEM of three independent experiments. ns = non-significant; * P < 0.05, ** P < 0.01; *** P < 0.001 (Student’s t-test).

Intriguingly, we noticed that the autophagosomes were localized close to the nucleus upon BHRF1 expression (***Figure 7D***). MT hyperacetylation is required for this juxtanuclear localization, suggesting its involvement in BHRF1-induced mitophagy. We therefore investigated the colocalization between mitochondria and autophagosomes in cells co-expressing α-tubulin K40A and BHRF1. When MT hyperacetylation was prevented, mitochondria did not colocalize anymore with autophagosomes (***Figure 7E***). The quantification of colocalization intensity, measured by Manders’ coefficient, confirmed that mitochondrial sequestration in autophagic vesicles did not occur when the mito-aggresome formation was impeded (***Figure 7F***). Altogether, this suggests that BHRF1-induced MT hyperacetylation triggers the localization of autophagosomes in the close vicinity of mitochondria.

### Autophagy is required for MT hyperacetylation

Our results demonstrated that the level of tubulin acetylation does not impact BHRF1-induced autophagy. We then decided to investigate whether, conversely, autophagy was involved in MT hyperacetylation. We first observed that inhibition of autophagy by treatment with 3-methyladenine (3-MA) or spautin-1 impeded the MT hyperacetylation (***Figure 8A***). To reinforce this finding, we used a genetic approach. We generated an ATG5 knockout (KO) cell line using a CRISPR/Cas9 approach, which is deficient for autophagy (***Figure 8B***). We observed that in ATG5 KO cells, BHRF1 was unable to increase the tubulin acetylation (***Figure 8C***). Moreover, mito-aggresome formation is dramatically decreased in autophagy-deficient cells (***Figures 8D and 8E***), in accordance with our previous results in presence of autophagy inhibitors (Vilmen *et al*., 2020). To confirm whether autophagy was also required during EBV reactivation, we treated Akata cells with spautin-1. We observed the lack of MT hyperacetylation (***Figure 8 – figure supplement 1A***) and of mito-aggresome formation (***Figure 8 – figure supplement 1B and 1C***) in EBV-reactivated treated cells.

**Figure 8.**
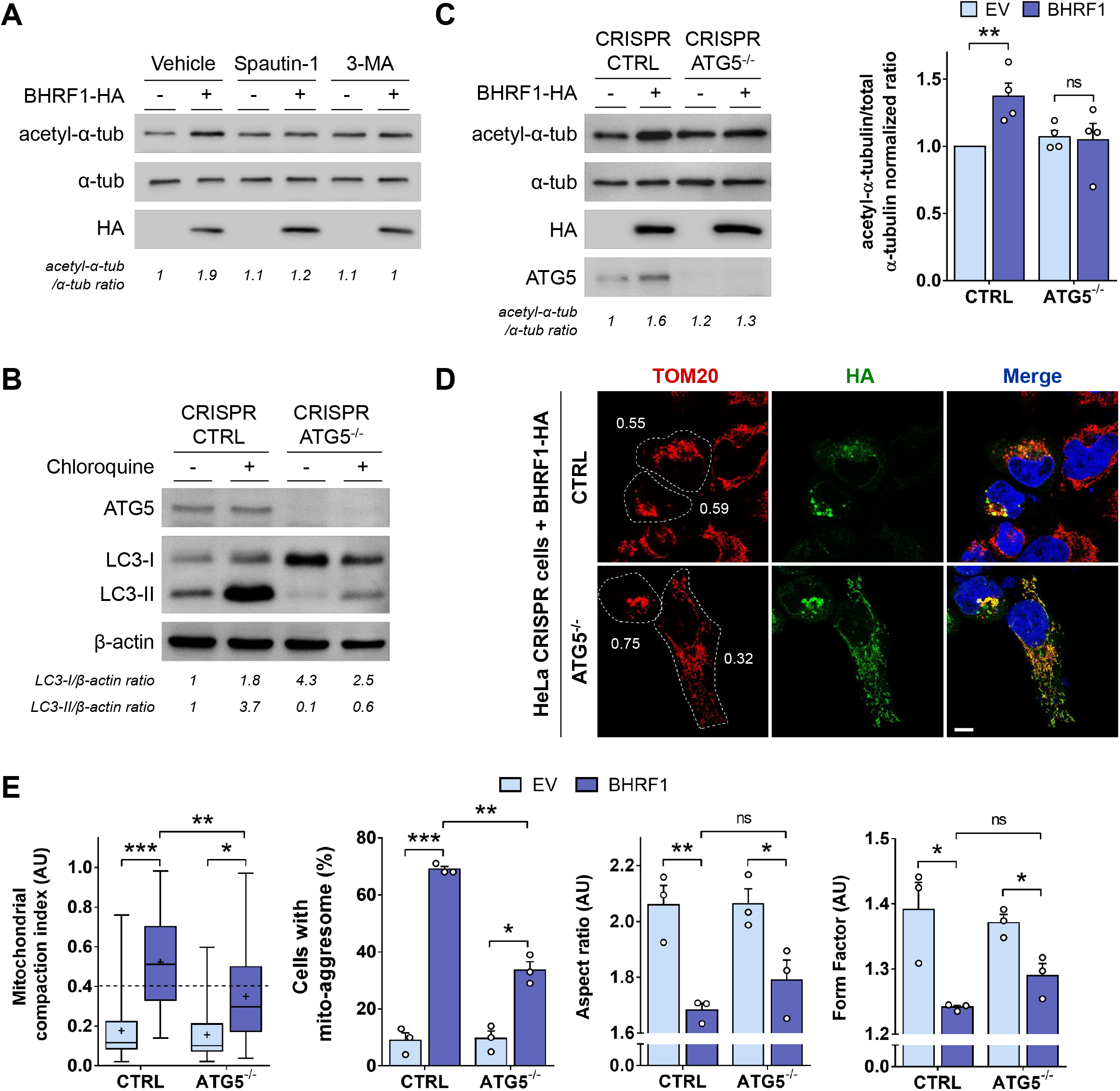
Autophagy inhibition prevents BHRF1-induced hyperacetylation and mito-aggresome formation. (**A**) Immunoblot analysis of acetyl-α-tubulin and α-tubulin of HeLa cells transfected with BHRF1-HA and treated or not with autophagic inhibitors (spautin-1 or 3-MA). (**B**) Control of the autophagy level and ATG5 extinction in HeLa cells treated or not with chloroquine. Immunoblot analysis of ATG5 and LC3. β-actin was used as a loading control. (**C-E**) HeLa cells deficient ATG5 (CRISPR ATG5-/-) and control cells (CRISPR CTRL) were transfected with BHRF1-HA for 24 h. (**C**) Left, immunoblot analysis of acetyl-α-tubulin, α-tubulin, HA and ATG5. Right, normalized ratios of acetyl-α-tubulin to α-tubulin. (**D**) Confocal images of cells immunostained for TOM20 and HA. Nuclei were stained with DAPI. Values of mitochondrial CI are indicated on representative cells. Scale bar: 10 μm. (**E**) Quantification of CI, percentage of cells with a mito-aggresome and mitochondrial fission parameters (n = 20 cells per condition). Data represent the mean ± SEM of three independent experiments. ns = non-significant; * P < 0.05; ** P < 0.01; *** P < 0.001 (Student’s t-test).

Taken together, our results demonstrated that BHRF1 needs autophagy activation to induce MT hyperacetylation. Therefore, we uncovered a novel mechanism to counteract antiviral immunity, involving the initiation of mitophagy through the induction of tubulin hyperacetylation by BHRF1.

## Discussion

Numerous strategies are used by viruses to counteract or evade the innate immune response. We describe here a novel mechanism for viral immune escape. We report for the first time that a post-translational modification of the cytoskeleton, MT hyperacetylation induced by EBV-encoded BHRF1, can block IFN production (***Figure 9***). We previously demonstrated that BHRF1 blocks the IFN response by modifying the mitochondrial dynamics and by stimulating autophagy (Vilmen *et al*., 2020). Expression of this viral protein induces mitochondrial fission, followed by the delocalization of fragmented mitochondria close to the nucleus in the proximity of the centrosome (***Figure 1 and Figure 1 – figure supplement 2***). Mitochondria are usually driven over long distances along MTs by motor proteins, thanks to the Miro/TRAK adaptor complex (Kruppa and Buss, 2021). In this study, we demonstrate that the MT network is hijacked by BHRF1 to ensure the mito-aggresome formation and to consequently block the antiviral innate immunity (***Figures 1 and 4***). We notably observed that disruption of the MT network impedes mitochondrial trafficking to the centrosome, suggesting a dynamic process. In accordance, our results indicate that BHRF1 requires the MT-based motor dynein, that typically moves towards the minus-end of MT, to regroup mitochondria in the juxtanuclear region (***Figure 6 and Figure 6 – figure supplement 1***).

**Figure 9.**
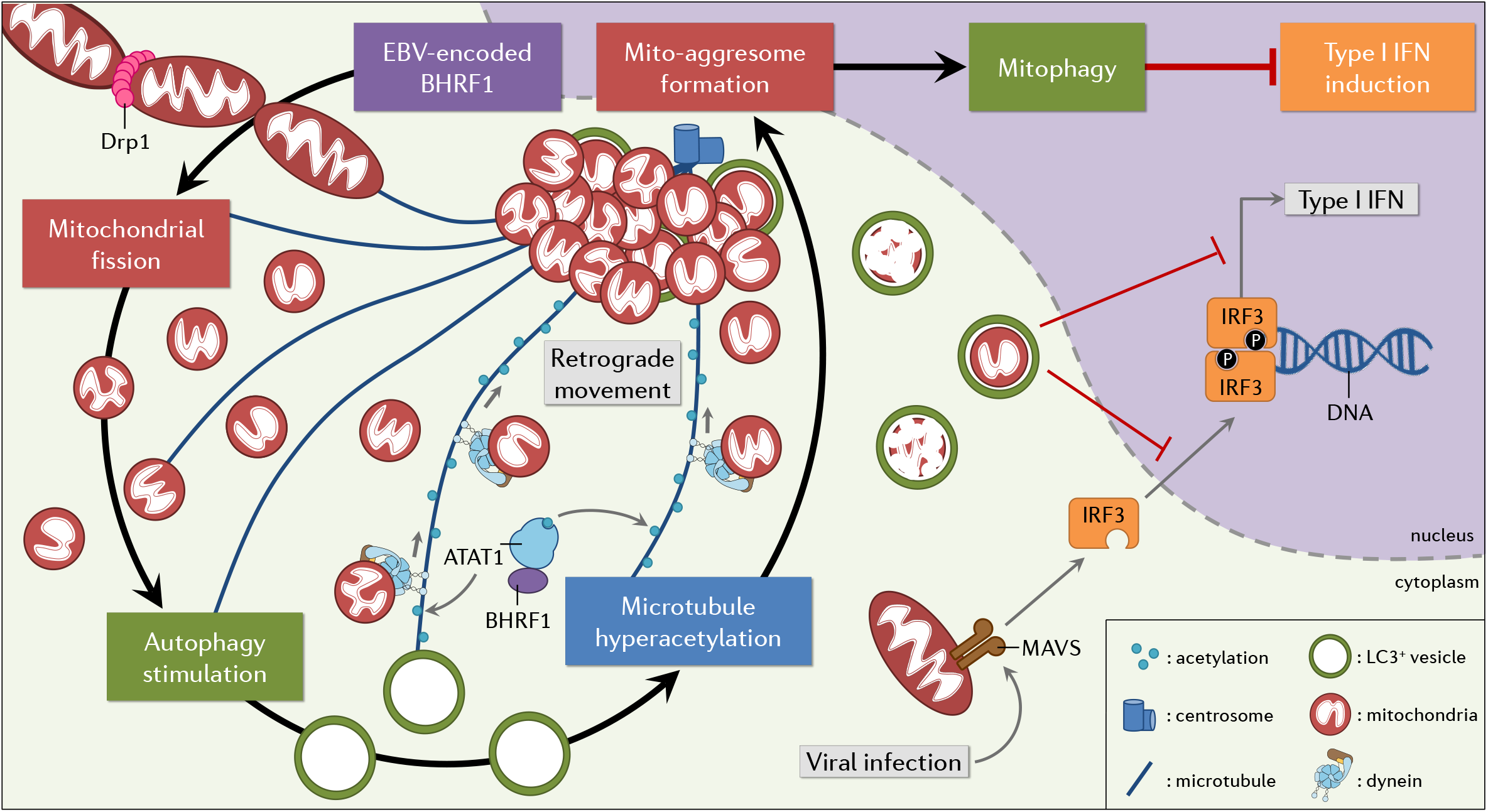
Working model. EBV encodes a Bcl-2 homolog that dampens type I IFN induction thanks to MT hyperacetylation. BHRF1 induces Drp1-mediated mitochondrial fission and subsequently autophagy. These cellular modifications lead to the mito-aggresome formation, and we demonstrated that the MT network and in particular its acetylation level are essential for this mitochondrial phenotype. Moreover, BHRF1 interacts with ATAT1 and induces MT hyperacetylation through ATAT1 activation. This hyperacetylation allows the retrograde transport of mitochondria in a dynein-dependent manner. Finally, sequestration and degradation of mitochondria block the innate immunity.

In eukaryotic cells, MTs are generally built from 13 chains of α-β tubulin dimers known as protofilaments. Properties and behaviors of MTs can be modulated by numerous post-translational modifications on both α and β tubulin (Janke and Magiera, 2020; Janke and Montagnac, 2017). We observed that the expression of BHRF1, after either transfection or EBV reactivation, increases the acetylation of α-tubulin on the lysine in position 40 (K40) (***Figure 2***). Moreover, acetylated MTs are required for the mito-aggresome formation and modulates the localization of autophagosomes (***Figures 3 and 7***). Acetylation of α-tubulin subtly changes its structure and disrupts the interaction between adjacent α-tubulins of neighboring protofilaments (Eshun-Wilson *et al*., 2019). The loss of this connection weakens interactions between protofilaments, which facilitates their sliding and decreases the flexural rigidity of microtubules, making them more resistant to mechanical bending-induced breakage and disassembly (Portran *et al*., 2017). Moreover, acetylated α-tubulin is more abundant in stable MTs, which are particularly implicated in mitochondrial transport (Friedman *et al*., 2010). Similar to what was observed with BHRF1, Misawa *et al*. demonstrated that accumulation of acetylated α-tubulin by KD of SIRT2 results in an aberrant mitochondrial distribution in the perinuclear region (Misawa *et al*., 2013).

Our results demonstrate that BHRF1 induces MT hyperacetylation by activating the acetyltransferase ATAT1. Indeed, BHRF1 interacts with ATAT1 and its KD by siRNA impedes the acetylation of α-tubulin and the mito-aggresome formation (***Figure 5***). Moreover, expression of the non-acetylatable K40A mutant form of tubulin confirmed that tubulin acetylation, and not just ATAT1 activity, is required for the mito-aggresome formation and sequestration of mitochondria in autophagosomes (***Figure 3***). We nevertheless investigated a potential inhibitory role of BHRF1 on tubulin deacetylases. However, our results showed that BHRF1 does not impact the global tubulin deacetylase activity, suggesting that HDAC6 and SIRT2, the two main α-tubulin deacetylases, are not inhibited by BHRF1 (Hubbert *et al*.,2002; North *et al*., 2003).

BHRF1-induced acetylation of α-tubulin could increase dynein recruitment to MTs, facilitating the traffic of fragmented mitochondria and autophagosomes to the centrosome. Indeed, it has been shown that MT hyperacetylation *in vitro* and in cells causes the recruitment of dynein and kinesin-1 to MTs, which notably stimulates anterograde and retrograde transport of organelles in neurons or epithelial cells (Dompierre *et al*., 2007; Geeraert *et al*., 2010). We know that BHRF1 interacts with BECN1, a key regulator of autophagosome formation and with BIM, a proapoptotic BH3-only protein expressed at the mitochondrial outer membrane (Desbien *et al*., 2009; Kvansakul *et al*., 2010; Vilmen *et al*.,2020). Moreover, it has been reported that BIM can also interact with the dynein light chain subunit DYNLL1 (Puthalakath *et al*., 1999; Singh *et al*., 2019). Interestingly, it has been shown that BIM binds to BECN1, linking BECN1 to dynein via DYNLL1 (Janke and Magiera, 2020; Luo *et al*., 2012; Roberts *et al*., 2013). Therefore, we hypothesize that BIM could connect mitochondria to MTs by interacting with BHRF1, BECN1 and DYNLL1. The formation of this putative complex could explain the mitochondrial movement toward the centrosome and participate in autophagy regulation.

It is interesting to note that MT hyperacetylation induced by BHRF1 or EBV reactivation is not involved in mitochondrial fission since expression of the non-acetylatable α-tubulin mutant K40A or the KD of ATAT1 have no impact on mitochondrial length or branching parameters (***Figures 3 and 5***). We previously demonstrated that BHRF1 increases mitochondrial fragmentation in a Drp1-dependent manner (Vilmen *et al*., 2020). Moreover, BHRF1 increases the dephosphorylation of Drp1 in position Ser637, which is then recruited to mitochondria to allow mitochondrial fission (Vilmen *et al*., 2020). In our context, the lack of direct involvement of acetylated MTs on mitochondrial fission could seem quite surprising, since it has been previously shown that MT hyperacetylation triggered by acute hyperosmotic stress enhanced mitochondrial fission (Perdiz *et al*., 2017). However, in the context of this acute stress, mitochondrial fission is associated with an increased level of Ser616 phosphorylation of Drp1, a different post-translational modification from the one observed upon BHRF1 expression. The implication of hyperacetylated-MT in the mitochondrial fission process may thus differ according to the type of cellular stress.

We observed that BHRF1-induced autophagy is necessary to increase MT acetylation since autophagy deficiency impedes MT acetylation (***Figure 8***). This could seem unexpected since several studies have demonstrated the importance of MT acetylation to induce autophagosome formation and maturation, and the direct correlation between MT acetylation levels and the autophagy flux (Esteves *et al*., 2019; Geeraert *et al*., 2010; Xie *et al*., 2010). For example, starvation-induced autophagy requires MT acetylation to recruit dynein and kinesin-1 on MTs and to allow JNK (c-Jun N-terminal kinase) activation, leading to BECN1 release (Geeraert *et al*., 2010). Very recently, it has been shown that p27, a tumor suppressor which promotes autophagy in response to glucose starvation, stimulates MT acetylation by binding to and stabilizing ATAT1, to allow the delivery of autophagosomes in the proximity of the centrosome for efficient fusion with lysosomes (Nowosad *et al*., 2021). However, MTs are not involved in autophagosome formation in basal autophagy (Mackeh *et al*., 2013). Similarly, we demonstrated that BHRF1-induced MT hyperacetylation has no role in non-selective autophagy. However, we observed that it is essential to induce mitophagy since, in the absence of α-tubulin acetylation, BHRF1 does not induce mitochondrial sequestration into autophagosomes (***Figure 7***). This suggests that the formation of the autophagosomal membranes around mitochondria requires acetylated MTs.

Several viral proteins are known to control type I IFN response by stimulating mitophagy (Ding *et al*., 2017; Wang *et al*., 2019, 2020). The viral glycoprotein (Gn) encoded by Hantaan virus translocates to mitochondria, induces complete mitophagy which leads to MAVS degradation and subsequently impairs innate immune response (Wang *et al*., 2020). PB1-F2 is a non-structural protein encoded by influenza A virus and constitutively localized to the mitochondrial inner membrane space (Yoshizumi *et al*., 2014). At mitochondria, the interaction of PB1-F2 with LC3B and mitochondrial TUFM (Tu translation elongation factor) similarly leads to mitophagy induction, MAVS degradation and restrains type I IFN production (Wang *et al*., 2020). Whether these pro-viral mechanisms depend on the tubulin acetylation level needs to be investigated. Interestingly, we explored the impact of non-viral mitophagy inducers, such as AMBRA1-ActA, CCCP and O/A on MT hyperacetylation and mitochondrial morphodynamics. We demonstrated that they can block IFN production via mitophagy induction, whereas none of them modulated MT acetylation (***Figure 4 – figure supplement 1***). These results clearly demonstrate BHRF1 blocks IFN production by an original mechanism, involving MT acetylation-dependent mitophagy.

To conclude, we demonstrated how BHRF1, which is expressed during the early stages of EBV infection and some latency programs associated with lymphomas (Fitzsimmons and Kelly, 2017), reduces innate immunity by manipulating the MT network. We propose a model (***Figure 9***) in which EBV exploits ATAT1-dependent acetylated MTs to modulate mitochondrial morphodynamics, thereby draining fragmented mitochondria toward the centrosome, segregating them from IFN signaling pathways in mitophagosomes to block IFN production. We previously showed that BHRF1 expression induces mitochondrial fission activation and subsequently autophagy stimulation (***Figure 9***). We report here that MT hyperacetylation is downstream of these events. Beyond EBV infection and latency, these findings identify for the first time a role for acetylated MTs in regulating IFN response and innate immunity against viruses.

## Materials and methods

### Antibodies

Primary antibodies used in this study included anti-α-tubulin (Sigma-Aldrich, T6199), anti-acetyl-α-tubulin (Sigma-Aldrich, T6793), anti-ATG5 (Sigma-Aldrich, A0731), anti-β-actin (Merck Millipore, MAB1501), anti-BHRF1 (Merck Millipore, MAB8188), anti-c-Myc (Sigma-Aldrich, M4439), anti-Drp1 (Cell Signaling Technology, D6C7), anti-Ea-D (Santa Cruz Biotechnology, sc-58121), anti-GFP (Roche, 11814460001), anti-GFP (Santa Cruz Biotechnology, sc-9996), anti-HA (Santa Cruz Biotechnology, sc-9082 and sc-7392), anti-IRF3 (Novus Biologicals, SD2062), anti-LC3 (MBL International, PM036 and M151-3B), anti-pericentrin (Sigma-Aldrich, HPA016820), anti-TOM20 (BD Biosciences, 612278), anti-TOM20 (Cell Signaling Technology, D8T4N), anti-ZEBRA (Santa Cruz Biotechnology, sc-53904).

Secondary antibodies used in this study included Alexa Fluor 488 goat anti-rabbit, anti-mouse (Jackson ImmunoResearch, 111-545-003, 115-545-003), Alexa Fluor 555 goat anti-rabbit, anti-mouse (Life technology, A21428, A21424), Alexa Fluor 647 donkey anti-rabbit, anti-mouse (Life technology, A31573, A31571). Horseradish peroxidase (HRP)-labeled goat anti-mouse and anti-rabbit secondary antibodies were purchased from Jackson ImmunoResearch Laboratories (115–035–003, 111–035–003).

### Cell culture and transfection

HeLa and HEK293T cells were cultured at 37°C under 5% CO_2_ in Dulbecco’s modified eagle medium (DMEM; Gibco, 41965-039) supplemented with 10% fetal calf serum (FCS; Dominique Dutscher, S181B-500). HeLa cells KD for Drp1 were previously described (Vilmen *et al*., 2020). ATG5 KO and control cell lines were generated from HeLa cells (see below) and cultured at 37°C under 5% CO_2_ in DMEM supplemented with 10% FCS and 1.25 μg/mL of puromycin. Akata cells were grown at 37°C under 5% CO_2_ in RPMI-1640 supplemented with 10% FCS and 2 mM of L-Glutamine (Gibco, 25030-081). Cells KD for BHRF1 were generated from Akata cells (see below) and cultured in the same media supplemented with 0.5 μg/mL of puromycin. Ramos cells were kindly provided by Joëlle Wiels (Institut Gustave Roussy, Villejuif, France) and grown in the same way as Akata cells. HEK293/EBV^+^ cells harboring WT and ΔBHRF1 bacmids were kindly provided by Wolfgang Hammerschmidt (German Research Center for Environmental Health, Munich, Germany) and were cultured at 37°C under 5% CO_2_ in DMEM supplemented with 10% FCS and 100 μg/mL of hygromycin B (Invitrogen, 10687010).

To reactivate EBV, Akata cells were treated with polyclonal rabbit anti-human IgG (Agilent, A0423) at 7.5 μg/mL for 8 or 24 h. To induce EBV productive cycle, HEK293/EBV WT and ΔBHRF1 mutant cells were transfected with ZEBRA and Rta expression plasmid (Altmann and Hammerschmidt, 2005).

DNA plasmid transfections were performed using Fugene HD transfection reagent (Promega Corporation, E2311) according to the manufacturer’s protocol. One day prior to transfection, cells were seeded in a 24-well plate and incubated at 37°C. On the day of transfection, DNA plasmids were dispersed in opti-MEM (Gibco, 31985-047) and Fugene HD was added to the mix (ratio DNA:Fugene equals 1:3). After incubation for 10 min at room temperature (RT), the transfection mix was added to the cells. Then cells were incubated at 37°C and 6 h post-transfection, DMEM supplemented with 10% FCS was added. Depending on the expression plasmids, cells were fixed 24 h or 48 h post-transfection.

### Co-immunoprecipitation assay

HeLa cells were cultured in a 6-well plate and transfected to co-express BHRF1-HA (or EV) and GFP-ATAT1 (or EV). One day after transfection, cells were washed with cold PBS and then lysed at 4°C in RIPA buffer (Thermo Fisher Scientific, 89900) supplemented with a cocktail of protease inhibitors. Then, extracts were centrifuged at 11’000xg at 4°C for 30 min to remove cell debris. Then, BHRF1 or ATAT1 was immunoprecipitated with a rabbit anti-HA antibody (2 μg) or with a mouse anti-GFP antibody (2 μg), respectively. Immunoprecipitations were performed overnight at 4°C. Protein A or G-Sepharose beads (Ademtech, 4230 or 4330) were added for 2 h at 4°C and were washed three times with RIPA buffer. The immune complexes were finally boiled for 10 min in loading buffer (see composition below) before being analyzed by SDS-PAGE.

### Dye and drugs

MitoTracker Red CMXRos was purchased from Life Technologies (M7512) and used at 100 nM for 30 min before cell fixation to stain the mitochondrial network. Phalloidin was purchased from Sigma-Aldrich (P1951) and used at 20 μg/mL for 1 h to stain F-actin. DAPI was purchased from Thermo Fisher Scientific (D1306) and used at 300 nM to stain dsDNA (nuclei).

Nocodazole was purchased from Sigma-Aldrich (M1404) and used at 10 μM to disrupt the MT network. Cytochalasin B was purchased from Sigma-Aldrich (C6762) and used at 10 μg/mL for 30 min to disrupt F-actin. Ciliobrevin D was purchased from Sigma-Aldrich (250401) and used at 40 μM overnight to inhibit cytosolic dyneins. To induce hyperosmotic shock, cells were cultured in media supplemented with 125 mM of NaCl for 30 min. Mdivi-1 was purchased from Sigma Aldrich (M0199) and used at 50 μM for 24 h to block Drp1 activity. Autophagic flux was monitored by the addition of CQ (Sigma-Aldrich, C6628) at 50 μM for 2 to 4 h. To inhibit autophagy, we used two different ways: cells were treated either with 3-MA (Sigma-Aldrich, M9281) at 5 mM for 8 h, or with spautin-1 (Sigma-Aldrich, SML0440) at 20 μM for 8 h. To induce mitophagy, cells were treated either during 4 h with CCCP (Sigma-Aldrich, C2759) at 10 μM, or during 6 h with a combination of oligomycin (Millipore, 495455) and antimycin A (Sigma-Aldrich, A8674) at 2.5 μM and 800 nM, respectively.

### Extinction of ATAT1 expression

To deplete ATAT1 in HeLa cells, we performed transfection of siRNA directed against ATAT1 encoding mRNA (or non-relevant siRNA as control). We used ON-TARGETplus siRNAs from Dharmacon as a pool of four siRNAs: 5-GUAGCUAGGUCCCGAUAUA-3; 5-GAGUAUAGCUAGAUCCCUU-3; 5-GGGAAACUCACC AGAACGA-3; and 5-CUUGUGAGAUUGUCGAGAU-3. The transfections of siRNA were performed using Oligofectamine reagent (Thermo Fisher Scientific; 12.252.011) according to the manufacturer’s protocol and were repeated at days 1, 2, and 3 at a final concentration of 100 nM. At the end of day 3, transfection with BHRF1-HA was also performed using Fugene HD transfection reagent as described above. Cells were then used at day 4 for immunoblotting or immunofluorescence analyses.

### Generation of Akata cells KD for BHRF1 expression

To knock down the expression of BHRF1 in Akata cells, we used an shRNA approach delivery through lentivirus transduction. Three different shRNA sequences directed against *BHRF1* mRNA were designed: 5’-CCGGGCCAGAGGACACTGTAGTTCTCTCGAGAGAACTACAGTGTCCTCTGGCTTTTTG-3’ (Fw, position 123), 5’-AATTCAAAAAGCCAGAGGACACTGTAGTTCTCTCGAGAGAACTACAGTGTCCTCTGGC-3’ (Rv, position 123), 5’-CCGGGTGTTGCTTGAGGAGATAATTCTCGAGAATTATCTCCTCAAGCAACACTTTTTG-3’ (Fw, position 154), 5’-AATTCAAAAAGTGTTGCTTGAGGAGATAATTCTCGAGAATTATCTCCTCAAGCAACAC-3’ (Rv, position 154), 5’-CCGGGGCGGCTGGTCTACATTAATTCTCGAGAATTAATGTAGACCAGCCGCCTTTTTG-3’ (Fw, position 442), 5’-AATTCAAAAAGGCGGCTGGTCTACATTAATTCTCGAGAATTAATGTAGACCAGCCGCC-3’ (Rv, position 442). These shRNA sequences were cloned into a pLK0.1-TRC vector, according to Addgene recommendations. To produce recombinant lentiviral particles, HEK293T cells (3×10^6^ cells) were transfected with 1.5 μg of pLK0.1-TRC/sh-BHRF1 plasmid, 1.125 μg of lentiviral packaging plasmid (psPAX2) and 375 ng of VSV-G envelope expressing plasmid (pMD2.G). Two days after transfection, the cell culture supernatant was collected and filtered (0.45 μm). The lentiviral particle stocks were directly used to transduce target cells by spinoculation method (centrifugation at 800xg at 32°C for 30 min). Three days after transduction, sh-BHRF1 expressing Akata cells were selected by addition of 0.5 μg/mL of puromycin in culture media.

### Generation of HeLa cells KO for ATG5 expression

Stock of lentiviral particles were obtained by transfection of HEK293T cells (2×10^6^ cells) with 2 μg of lentivector plasmid lentiCRISPRv2 (a gift from Feng Zhang, Addgene plasmid, 52961) expressing single guide RNA targeting an exon within ATG5 gene, using Lipofectamine LTX with PLUS Reagent (Life technologies), along with 3 μg of psPAX2 and 1 μg of pMD2.G. The following sequence was used to target ATG5 genes: 5’-AAGATGTGCTTCGAGATGTG-3’ (ATG5, exon 1; guide 92). The sequence 5’-GTTCCGCGTTACATAACTTA-3’ was used as negative control (referred as CTRL) and previously described (Kearns *et al*., 2014). Twenty-four hours after transfection, cell media were removed, and cells were cultured for additional 24 h in fresh media. Supernatants were then collected, filtered (0.45 μm) and used to transduce HeLa cells. Three days after transduction, cells were selected by addition of 1.25 μg/mL of puromycin in culture media.

### *In vitro* tubulin deacetylation assay

Cell lysates of control cells or cells transfected with BHRF1-HA plasmids were prepared using 10 mM Tris buffer pH 8.0, supplemented with 1% Triton, 1 mM NAD^+^, and the protease inhibitor cocktail. Lysate aliquots containing 20 μg of proteins were mixed with 2 μg of soluble acetylated tubulin dimers purified from porcine brain (Mackeh *et al*., 2014; Walker *et al*., 1988) and adjusted to 50 μL with NAD^+^-containing PEM buffer (80 mM PIPES, 2 mM EGTA, 1 mM MgCl_2_). After 1 h at 37°C, reactions were stopped by adding 4X SDS-PAGE sample buffer and immediately subjected to SDS-PAGE and Western blotting of acetylated and total α-tubulin.

### Immunoblot analysis

Cells were lysed in lysis buffer (65 mM Tris, pH 6.8, 4% SDS, 1.5%β-mercaptoethanol, and a cocktail of protease inhibitors) and kept at 100°C for 10 min. Protein extracts were resolved on SDS-PAGE (12.5%) and electro-transferred onto a polyvinylidene difluoride membrane (Amersham, 10600002). After 1h of incubation in blocking buffer (PBS, 0.1% Tween20, and 5% BSA or nonfat dry milk), the membranes were probed overnight at 4°C with primary antibodies. Horseradish peroxidase-labeled antibodies were used as secondary antibody, and revelation was performed using the ECL detection system according to the manufacturer’s instructions (Merck Millipore, WBKLS0500). Scanning for quantification of protein levels was monitored using ImageJ software. An anti-β-actin antibody was used to ensure equal loadings and normalize quantification.

### Immunofluorescence analysis

For immunofluorescence, adherent cells were cultured on glass coverslips in 24-well plates. The cell monolayers were washed with PBS and cells were fixed with paraformaldehyde (PFA; 4%) in PBS or methanol, depending on the antibodies. The cells were treated with PBS containing NH_4_Cl (50 mM) for 10 min, permeabilized using PBS, Triton X-100 (0.2%) for 4 min, washed twice with PBS and then incubated for 1 h in PBS, gelatin (0.2%) supplemented with FCS (5%) for blocking. Then the cells were incubated for 1 h with the appropriate primary antibody diluted in PBS, gelatin (0.2%) at RT. Cells were washed three times with PBS and then incubated with the appropriate secondary antibody diluted in PBS, gelatin (0.2%) for 1 h at RT. After washing, the nuclei were counterstained with DAPI. Coverslips were mounted in Glycergel (Agilent Dako, C056330-2) and observed using a Nikon Eclipse 80i epifluorescence microscope (Nikon Instruments) or a Leica TCS SP8 X inverted confocal microscope (Leica, USA). Photographic images were resized, organized, and labeled using Fiji-ImageJ software or LAS AF Lite. Imaris software was also used to perform 3D reconstruction videos.

A specific procedure was performed for the colocalization study between mitochondria and autophagosomes. Permeabilization and blocking were performed by incubating the cells for 30 min with PBS, saponin (0.075%; Sigma-Aldrich, 84,510), BSA (1%), and FCS (5%). Antibodies were diluted in PBS, saponin (0.075%), and BSA (1%). In addition, between each antibody incubation, cells were washed twice in PBS, saponin (0.075%), BSA (1%), once with PBS, saponin (0.037%), BSA (0.5%) and twice in PBS. Other steps were the same as described above.

To immunostain suspension cells, they were washed once with PBS and diluted at the concentration of 10^6^ cells/mL. Then, 15 μL of this cell suspension were dropped on a slide and dried for 10 min at 37°C. A circle was drawn around the cell footprint with a hydrophobic PAP pen. Cells were then fixed with PFA (4%), and the next steps were the same as described above.

### Luciferase reporter assays to assess IFN-β promoter activity

HEK293T cells were cultured in 24-well plates and transfected for 24 h using Fugene HD, with plasmids encoding the IFN-β luciferase reporter (firefly luciferase; 100 ng), pRL-TK (renilla luciferase; 25 ng), pΔRIG-I (2xCARD) (encoding a positive dominant form of RIG-I; 150 ng), plasmids expressing BHRF1-HA (or EV as control; 150 ng) and mCherry-Tub (WT or K40A; 150 ng). One day after transfection, cells were lysed and measurement of firefly and renilla luciferase activities was performed using the dual-luciferase reporter assay system (Promega Corporation, E1910) according to the manufacturer’s protocol. Relative expression levels were calculated by dividing the firefly luciferase values by those of renilla luciferase and normalized to the control condition (without BHRF1-HA expression).

### Microtubule depolymerization and repolymerization

To depolymerize MTs, HeLa cells were seeded directly onto glass coverslips and incubated one day before being transfected to express BHRF1. Twenty-four hours after transfection, cells were treated with 10 μM nocodazole (Sigma-Aldrich, M1404) for 2 h at 37°C then 1.5 h at 4°C, to depolymerize the MT network. Cells were immediately PFA 4% fixed and then methanol-permeabilized (t = 0 h before repolymerization). MTs and mitochondria were visualized by immunostaining of α-tubulin and TOM20, respectively. To allow reassembly of the MT network, nocodazole was removed by repeated washes with PBS, and cells were left 1 h in warm complete medium (t = 1 h after MTs regrowth). Cells were then immunostained for BHRF1, TOM20, and α-tubulin.

### Mitochondrial phenotype assessment

BHRF1-induced mito-aggresomes were scored by measuring the CI of mitochondrial staining, as we previously described (Vilmen *et al*., 2020), based on Narendra et al. study (Narendra *et al*., 2010). We arbitrary determined that a cell presents a mito-aggresome when the CI is above 0.4. The percentage of cells presenting a mito-aggresome was determined by counting at least 20 random cells in each condition from three independent experiments.

Mitochondrial morphology was assessed by measuring AR and FF, which are indicators of mitochondrial length and branching, respectively. These parameters were quantified as described before (Koopman *et al*., 2006). Briefly, on Fiji-ImageJ software, images of mitochondria were convolved using the matrix developed by Koopman, then a threshold was applied to isolate mitochondria from the background. Then cell outline was drawn and mitochondrial phenotype was assessed by the implement analysis particle tool. The AR value is given by the software and corresponds to the ratio between the major and minor axis of the ellipsis equivalent to the mitochondria. The FF value is defined as (P_mit_^2^)/(4πA_mit_), where Pmit is the length of mitochondrial outline and A_mit_ is the area of mitochondria. The mitochondrial fission parameters were determined by analyzing at least 15 random cells in each condition from three independent experiments.

### Plasmids

We previously described the BHRF1-HA expression vector (Vilmen *et al*., 2020), which was constructed from a pcDNA3.1 expression vector (EV) purchased from Invitrogen (V79020). The mCherry-α-tubulin WT-K40K control vector was kindly provided by Dr. R. Y. Tsien (Howard Hughes Medical Center, University of San Diego, USA) (Shaner *et al*.,2004) and the mutated vector (mCherry-α-tubulin K40A) was a kind gift from Dr. F. Saudou (Institut Curie, Orsay, France) (Dompierre *et al*., 2007). The GFP-ATAT1 expression plasmid (27099) and the pLK0.1-TRC cloning vector (10878) were purchased from Addgene. The plasmids Flag-ΔRIG-I (2xCARD) and pIFN-β-Luc were previously described (Yoneyama *et al*., 2004, 1996) and kindly provided by Dr. C. Lagaudrière (I2BC, Gif-sur-Yvette, France). The plasmid pRL-TK was purchased from Promega Corporation (E2241). The p50-dynamitin construct was a gift from Dr. S. Etienne-Manneville (Institut Pasteur, Paris, France) (Etienne-Manneville and Hall, 2001). ZEBRA and Rta expression plasmids were kindly provided by Henri Gruffat (CIRI, Lyon, France). The AMBRA1-ActA expression plasmid was previously described (Strappazzon *et al*., 2015) and a gift from Flavie Strappazzon. The plasmids psPAX2 (Addgene, 12260) and pMD2.G (Addgene, 12259) come from Didier Trono lab (EPFL, Lausanne, Switzerland).

### Results quantification

The number of endogenous LC3 dots per cell was determined using Fiji-ImageJ software. All images were taken with the same microscope settings to allow for comparison. Images were converted into 8-bit. Cells’ outlines were drawn and nuclei cropped to conserve only the cytoplasm. Thresholding of images was performed to detect only LC3 vesicles. Note that the value of the threshold was conserved and used for each analyzed cell. Finally, the “analyze particles” plug-in was used to count the total number of LC3 dots per cell. For each condition, at least 25 cells were analyzed from three independent experiments.

The distance between autophagosomes and the nucleus was measured using Fiji-ImageJ software. For each cell, the distance of all autophagosomes to the nucleus was measured and the mean distance was considered for the statistical analysis. For each condition, at least 20 random cells were analyzed from three independent experiments.

The colocalization degree between mitochondrial staining and autophagosomes staining was assessed by measuring Manders split coefficient using the “colocalization threshold” plug-in on Fiji-ImageJ software. For each condition, 30 random cells were analyzed.

The proportion of cells presenting an IRF3 nuclear localization was determined by counting at least 50 random cells in each condition from three independent experiments.

### Statistical analysis

Data are expressed as mean ± standard error of the mean (SEM) and were analyzed with Prism software (GraphPad) by using Student’s t-test. P values less than 0.05 were considered statistically significant. Data from three independent experiments were analyzed.

## Acknowledgments

We would like to thank for her technical assistance Valérie Nicolas (Université Paris-Saclay, Châtenay-Malabry, France) from the MIPSIT cellular imaging facility. Also, the present work has benefited from the core facilities of Imagerie-Gif, (http://www.i2bc.paris-saclay.fr), member of IBiSA (http://www.ibisa.net), supported by “France-Biolmaging” (ANR-10-INBS-04-01), and the Labex “Saclay Plant Science” (ANR-11-IDEX-0003-02). We wish to thank Flavie Strappazzon for providing us AMBRA1-ActA plasmid and we are also very grateful to Joëlle Wiels for giving us Ramos cells. Finally, we thank Stephanie Straubel for critical reading of the manuscript.

## Additional information

### Funding

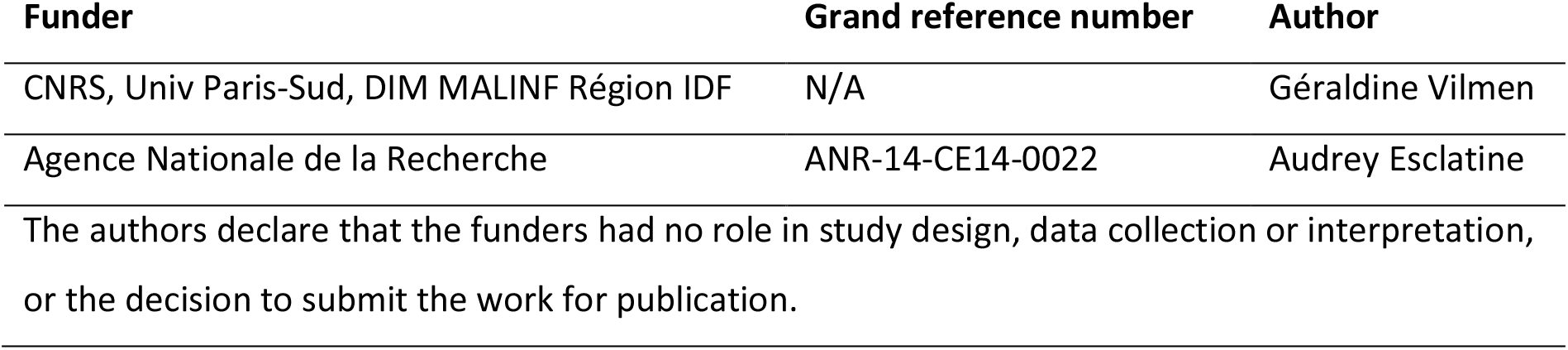

### Author contributions

DG, GV, DP, GB, conception and design, investigation, formal analysis, writing original draft, writing review and editing; EH, investigation, formal analysis; VM, formal analysis, funding acquisition, supervision; CP, FQ, CBT, formal analysis, provided materials; ML, conception and design, formal analysis, writing original draft, supervision; AE, conception and design, formal analysis, funding acquisition, writing original draft, writing review and editing, supervision.

## Additional files

### Figure supplements

- Figure 1 – figure supplement 1. Knockdown of BHRF1 expression in Akata cells prevents mito-aggresome formation after EBV reactivation.
- Figure 1 – figure supplement 2. BHRF1 localizes in the proximity of the centrosome.
- Figure 1 – figure supplement 3. The actin network is not involved in the formation of BHRF1 induced mito-aggresomes.
- Figure 3 – figure supplement 1. Impact of the non-acetylatable α-tubulin expression.
- Figure 3 – figure supplement 2. MT hyperacetylation required mitochondrial fission.
- Figure 4 – figure supplement 1. Interplay between MT hyperacetylation and IFN inhibition in response to different mitophagy inducers.
- Figure 5 – figure supplement 1. Loss of ATAT1 has no impact on mitochondrial network.
- Figure 5 – figure supplement 2. ATAT1 colocalizes with BHRF1.
- Figure 6 – figure supplement 1. Dynein inhibition prevents BHRF1-induced mito-aggresomes.
- Figure 7 – figure supplement 1. BHRF1 stimulates autophagy when α-tubulin WT is expressed.
- Figure 8 – figure supplement 1. Autophagy inhibition prevents MT hyperacetylation and mito-aggresome formation during EBV reactivation.

### Data availability

All data generated or analyzed during this study are included in this published article and its supplementary information files.

## Notes

### Competing Interest Statement

The authors have declared no competing interest.

## References

Altmann M, Hammerschmidt W. 2005. Epstein-Barr Virus Provides a New Paradigm: A Requirement for the Immediate Inhibition of Apoptosis. PLoS Biol 3:e404. doi:10.1371/journal.pbio.0030404

Arnoult D, Soares F, Tattoli I, Girardin SE. 2011. Mitochondria in innate immunity. EMBO Rep 12:901–910. doi:10.1038/embor.2011.157

Castanier C, Garcin D, Vazquez A, Arnoult D. 2010. Mitochondrial dynamics regulate the RIG-I-like receptor antiviral pathway. EMBO Rep 11:133–138. doi:10.1038/embor.2009.258

Chatel-Chaix L, Cortese M, Romero-Brey I, Bender S, Neufeldt CJ, Fischl W, Scaturro P, Schieber N, Schwab Y, Fischer B, Ruggieri A, Bartenschlager R. 2016. Dengue Virus Perturbs Mitochondrial Morphodynamics to Dampen Innate Immune Responses. Cell Host Microbe 20:342–356. doi:10.1016/j.chom.2016.07.008

De Vos KJ, Allan VJ, Grierson AJ, Sheetz MP. 2005. Mitochondrial Function and Actin Regulate Dynamin-Related Protein 1-Dependent Mitochondrial Fission. Curr Biol 15:678–683. doi:10.1016/j.cub.2005.02.064

Desbien AL, Kappler JW, Marrack P. 2009. The Epstein-Barr virus Bcl-2 homolog, BHRF1, blocks apoptosis by binding to a limited amount of Bim. Proc Natl Acad Sci U S A 106:5663–5668. doi:10.1073/pnas.0901036106

Detmer SA, Chan DC. 2007. Functions and dysfunctions of mitochondrial dynamics. Nat Rev Mol Cell Biol 8:870–879. doi:10.1038/nrm2275

Ding B, Zhang L, Li Z, Zhong Y, Tang Q, Qin Y, Chen M. 2017. The Matrix Protein of Human Parainfluenza Virus Type 3 Induces Mitophagy that Suppresses Interferon Responses. Cell Host & Microbe 21:538–547.e4. doi:10.1016/j.chom.2017.03.004

Dompierre JP, Godin JD, Charrin BC, Cordelieres FP, King SJ, Humbert S, Saudou F. 2007. Histone Deacetylase 6 Inhibition Compensates for the Transport Deficit in Huntington’s Disease by Increasing Tubulin Acetylation. J Neurosci 27:3571–3583. doi:10.1523/JNEUROSCI.0037-07.2007

Eshun-Wilson L, Zhang R, Portran D, Nachury MV, Toso DB, Löhr T, Vendruscolo M, Bonomi M, Fraser JS, Nogales E. 2019. Effects of α-tubulin acetylation on microtubule structure and stability. Proc Natl Acad Sci U S A 116:10366–10371. doi:10.1073/pnas.1900441116

Esteves AR, Palma AM, Gomes R, Santos D, Silva DF, Cardoso SM. 2019. Acetylation as a major determinant to microtubule-dependent autophagy: Relevance to Alzheimer’s and Parkinson disease pathology. Biochim Biophys Acta, Post-Translational Modifications In Brain Health And Disease 1865:2008–2023. doi:10.1016/j.bbadis.2018.11.014

Etienne-Manneville S, Hall A. 2001. Integrin-mediated activation of Cdc42 controls cell polarity in migrating astrocytes through PKCzeta. Cell 106:489–498. doi:10.1016/s0092-8674(01)00471-8

Fitzsimmons L, Kelly G. 2017. EBV and Apoptosis: The Viral Master Regulator of Cell Fate? Viruses 9:339–373. doi:10.3390/v9110339

Fransson Å, Ruusala A, Aspenström P. 2006. The atypical Rho GTPases Miro-1 and Miro-2 have essential roles in mitochondrial trafficking. Biochem Biophys Res Commun 344:500–510. doi:10.1016/j.bbrc.2006.03.163

Frederick RL, Shaw JM. 2007. Moving Mitochondria: Establishing Distribution of an Essential Organelle. Traffic 8:1668–1675. doi:10.1111/j.1600-0854.2007.00644.x

Friedman JR, Webster BM, Mastronarde DN, Verhey KJ, Voeltz GK. 2010. ER sliding dynamics and ER– mitochondrial contacts occur on acetylated microtubules. J Cell Biol 190:363–375. doi:10.1083/jcb.200911024

Geeraert C, Ratier A, Pfisterer SG, Perdiz D, Cantaloube I, Rouault A, Pattingre S, Proikas-Cezanne T, Codogno P, Poüs C. 2010. Starvation-induced Hyperacetylation of Tubulin Is Required for the Stimulation of Autophagy by Nutrient Deprivation. J Biol Chem 285:24184–24194. doi:10.1074/jbc.M109.091553

Green DR, Levine B. 2014. To Be or Not to Be? How Selective Autophagy and Cell Death Govern Cell Fate. Cell 157:65–75. doi:10.1016/j.cell.2014.02.049

Hancock WO. 2014. Bidirectional cargo transport: moving beyond tug of war. Nat Rev Mol Cell Biol 15:615–628. doi:10.1038/nrm3853

Hubbert C, Guardiola A, Shao R, Kawaguchi Y, Ito A, Nixon A, Yoshida M, Wang X-F, Yao T-P. 2002. HDAC6 is a microtubule-associated deacetylase. Nature 417:455–458. doi:10.1038/417455a

Janke C, Magiera MM. 2020. The tubulin code and its role in controlling microtubule properties and functions. Nat Rev Mol Cell Biol 21:307–326. doi:10.1038/s41580-020-0214-3

Janke C, Montagnac G. 2017. Causes and Consequences of Microtubule Acetylation. Curr Biol 27:1287–1292. doi:10.1016/j.cub.2017.10.044

Kaufmann T, Kukolj E, Brachner A, Beltzung E, Bruno M, Kostrhon S, Opravil S, Hudecz O, Mechtler K, Warren G, Slade D. 2016. SIRT2 regulates nuclear envelope reassembly through ANKLE2 deacetylation. J Cell Sci 129:4607–4621. doi:10.1242/jcs.192633

Kearns NA, Genga RMJ, Enuameh MS, Garber M, Wolfe SA, Maehr R. 2014. Cas9 effector-mediated regulation of transcription and differentiation in human pluripotent stem cells. Development 141:219–223. doi:10.1242/dev.103341

Khanim F, Mackett M, Young LS, Dawson J, Meseda CA, Dawson C. 1997. BHRF1, a viral homologue of the Bcl-2 oncogene, is conserved at both the sequence and functional level in different Epstein-Barr virus isolates. J Gen Virol 78:2987–2999. doi:10.1099/0022-1317-78-11-2987

Köchl R, Hu XW, Chan EYW, Tooze SA. 2006. Microtubules Facilitate Autophagosome Formation and Fusion of Autophagosomes with Endosomes: Role of Microtubules in AV Formation. Traffic 7:129–145. doi:10.1111/j.1600-0854.2005.00368.x

Koopman WJH, Visch H-J, Smeitink JAM, Willems PHGM. 2006. Simultaneous quantitative measurement and automated analysis of mitochondrial morphology, mass, potential, and motility in living human skin fibroblasts. Cytometry Part A 69A:1–12. doi:10.1002/cyto.a.20198

Kruppa AJ, Buss F. 2021. Motor proteins at the mitochondria-cytoskeleton interface. J Cell Sci 134:1–13. doi:10.1242/jcs.226084

Kvansakul M, Wei AH, Fletcher JI, Willis SN, Chen L, Roberts AW, Huang DCS, Colman PM. 2010. Structural Basis for Apoptosis Inhibition by Epstein-Barr Virus BHRF1. PLoS Pathog 6:e1001236. doi:10.1371/journal.ppat.1001236

Lazarou M, Sliter DA, Kane LA, Sarraf SA, Wang C, Burman JL, Sideris DP, Fogel AI, Youle RJ. 2015. The ubiquitin kinase PINK1 recruits autophagy receptors to induce mitophagy. Nature 524:309–314. doi:10.1038/nature14893

Lee AJ, Ashkar AA. 2018. The Dual Nature of Type I and Type II Interferons. Front Immunol 9:2061–2070. doi:10.3389/fimmu.2018.02061

López-Doménech G, Covill-Cooke C, Ivankovic D, Halff EF, Sheehan DF, Norkett R, Birsa N, Kittler JT. 2018. Miro proteins coordinate microtubule- and actin-dependent mitochondrial transport and distribution. EMBO J 37:321–336. doi:10.15252/embj.201696380

Luo S, Garcia-Arencibia M, Zhao R, Puri C, Toh PPC, Sadiq O, Rubinsztein DC. 2012. Bim inhibits autophagy by recruiting Beclin 1 to microtubules. Mol Cell 47:359–370. doi:10.1016/j.molcel.2012.05.040

Mackeh R, Lorin S, Ratier A, Mejdoubi-Charef N, Baillet A, Bruneel A, Hamaï A, Codogno P, Poüs C, Perdiz D. 2014. Reactive Oxygen Species, AMP-activated Protein Kinase, and the Transcription Cofactor p300 Regulate α-Tubulin Acetyltransferase-1 (αTAT-1/MEC-17)-dependent Microtubule Hyperacetylation during Cell Stress. J Biol Chem 289:11816–11828. doi:10.1074/jbc.M113.507400

Mackeh R, Perdiz D, Lorin S, Codogno P, Pous C. 2013. Autophagy and microtubules - new story, old players. J Cell Sci 126:1071–80. doi:10.1242/jcs.115626

Misawa T, Takahama M, Kozaki T, Lee H, Zou J, Saitoh T, Akira S. 2013. Microtubule-driven spatial arrangement of mitochondria promotes activation of the NLRP3 inflammasome. Nat Immunol 14:454–460. doi:10.1038/ni.2550

Narendra D, Kane LA, Hauser DN, Fearnley IM, Youle RJ. 2010. p62/SQSTM1 is required for Parkin-induced mitochondrial clustering but not mitophagy; VDAC1 is dispensable for both. Autophagy 6:1090–1106. doi:10.4161/auto.6.8.13426

Narendra DP, Tanaka A, Suen D-F, Youle RJ. 2008. Parkin is recruited selectively to impaired mitochondria and promotes their autophagy. J Cell Biol 183:795–803. doi:10.1083/jcb.200809125

Nowosad A, Creff J, Jeannot P, Culerrier R, Codogno P, Manenti S, Nguyen L, Besson A. 2021. p27 controls autophagic vesicle trafficking in glucose-deprived cells via the regulation of ATAT1-mediated microtubule acetylation. Cell Death Dis 12:1–18. doi:10.1038/s41419-021-03759-9

Perdiz D, Lorin S, Leroy-Gori I, Poüs C. 2017. Stress-induced hyperacetylation of microtubule enhances mitochondrial fission and modulates the phosphorylation of Drp1 at 616 Ser. Cell Signalling 39:32–43. doi:10.1016/j.cellsig.2017.07.020

Perdiz D, Mackeh R, Poüs C, Baillet A. 2011. The ins and outs of tubulin acetylation: More than just a post-translational modification? Cell Signalling 23:763–771. doi:10.1016/j.cellsig.2010.10.014

Portran D, Schaedel L, Xu Z, Théry M, Nachury MV. 2017. Tubulin acetylation protects long-lived microtubules against mechanical ageing. Nat Cell Biol 19:391–398. doi:10.1038/ncb3481

Puthalakath H, Huang DCS, O’Reilly LA, King SM, Strasser A. 1999. The Proapoptotic Activity of the Bcl-2 Family Member Bim Is Regulated by Interaction with the Dynein Motor Complex. Mol Cell 3:287–296. doi:10.1016/S1097-2765(00)80456-6

Reed NA, Cai D, Blasius TL, Jih GT, Meyhofer E, Gaertig J, Verhey KJ. 2006. Microtubule Acetylation Promotes Kinesin-1 Binding and Transport. Curr Biol 16:2166–2172. doi:10.1016/j.cub.2006.09.014

Roberts AJ, Kon T, Knight PJ, Sutoh K, Burgess SA. 2013. Functions and mechanics of dynein motor proteins. Nat Rev Mol Cell Biol 14:713–726. doi:10.1038/nrm3667

Shaner NC, Campbell RE, Steinbach PA, Giepmans BNG, Palmer AE, Tsien RY. 2004. Improved monomeric red, orange and yellow fluorescent proteins derived from Discosoma sp. red fluorescent protein. Nat Biotechnol 22:1567–1572. doi:10.1038/nbt1037

Shida T, Cueva JG, Xu Z, Goodman MB, Nachury MV. 2010. The major α-tubulin K40 acetyltransferase αTAT1 promotes rapid ciliogenesis and efficient mechanosensation. Proc Natl Acad Sci U S A 107:21517–21522. doi:10.1073/pnas.1013728107

Singh PK, Weber A, Häcker G. 2018. The established and the predicted roles of dynein light chain in the regulation of mitochondrial apoptosis. Cell Cycle 17:1037–1047. doi:10.1080/15384101.2018.1464851

Strappazzon F, Nazio F, Corrado M, Cianfanelli V, Romagnoli A, Fimia GM, Campello S, Nardacci R, Piacentini M, Campanella M, Cecconi F. 2015. AMBRA1 is able to induce mitophagy via LC3 binding, regardless of PARKIN and p62/SQSTM1. Cell Death Differ 22:419–32. doi:10.1038/cdd.2014.139

Tang B. 2018. Miro—Working beyond Mitochondria and Microtubules. Cells 7:18–24. doi:10.3390/cells7030018

Vilmen G, Glon D, Siracusano G, Lussignol M, Shao Z, Hernandez E, Perdiz D, Quignon F, Mouna L, Poüs C, Gruffat H, Maréchal V, Esclatine A. 2020. BHRF1, a BCL2 viral homolog, disturbs mitochondrial dynamics and stimulates mitophagy to dampen type I IFN induction. Autophagy 17:1296–1315. doi:10.1080/15548627.2020.1758416

Walker RA, O’Brien E. T, Pryer NK, Soboeiro MF, Voter WA, Erickson HP, Salmon ED. 1988. Dynamic instability of individual microtubules analyzed by video light microscopy: rate constants and transition frequencies. J Cell Biol 107:1437–1448. doi:10.1083/jcb.107.4.1437

Wang K, Ma H, Liu H, Ye W, Li Z, Cheng L, Zhang L, Lei Y, Shen L, Zhang F. 2019. The Glycoprotein and Nucleocapsid Protein of Hantaviruses Manipulate Autophagy Flux to Restrain Host Innate Immune Responses. Cell Reports 27:2075–2091.e5. doi:10.1016/j.celrep.2019.04.061

Wang R, Zhu Y, Ren C, Yang S, Tian S, Chen H, Jin M, Zhou H. 2020. Influenza A virus protein PB1-F2 impairs innate immunity by inducing mitophagy. Autophagy 1–16. doi:10.1080/15548627.2020.1725375

West AP, Shadel GS, Ghosh S. 2011. Mitochondria in innate immune responses. Nat Rev Immunol 11:389–402. doi:10.1038/nri2975

Xie R, Nguyen S, McKeehan WL, Liu L. 2010. Acetylated microtubules are required for fusion of autophagosomes with lysosomes. BMC Cell Biol 11:89–100. doi:10.1186/1471-2121-11-89

Yoneyama M, Kikuchi M, Natsukawa T, Shinobu N, Imaizumi T, Miyagishi M, Taira K, Akira S, Fujita T. 2004. The RNA helicase RIG-I has an essential function in double-stranded RNA-induced innate antiviral responses. Nat Immunol 5:730–737. doi:10.1038/ni1087

Yoneyama M, Suhara W, Fukuhara Y, Sato M, Ozato K, Fujita T. 1996. Autocrine Amplification of Type I Interferon Gene Expression Mediated by Interferon Stimulated Gene Factor 3 (ISGF3). J Biochem 120:160–169. doi:10.1093/oxfordjournals.jbchem.a021379

Yoshizumi T, Ichinohe T, Sasaki O, Otera H, Kawabata S, Mihara K, Koshiba T. 2014. Influenza A virus protein PB1-F2 translocates into mitochondria via Tom40 channels and impairs innate immunity. Nat Commun 5:4713. doi:10.1038/ncomms5713

Zheng K, Jiang Y, He Z, Kitazato K, Wang Y. 2017. Cellular defence or viral assist: the dilemma of HDAC6. J Gen Virol 98:322–337. doi:10.1099/jgv.0.000679

