## supplemental figures for "Essential role of hyperacetylated microtubules in innate immunity escape orchestrated by the EBV-encoded BHRF1 protein"

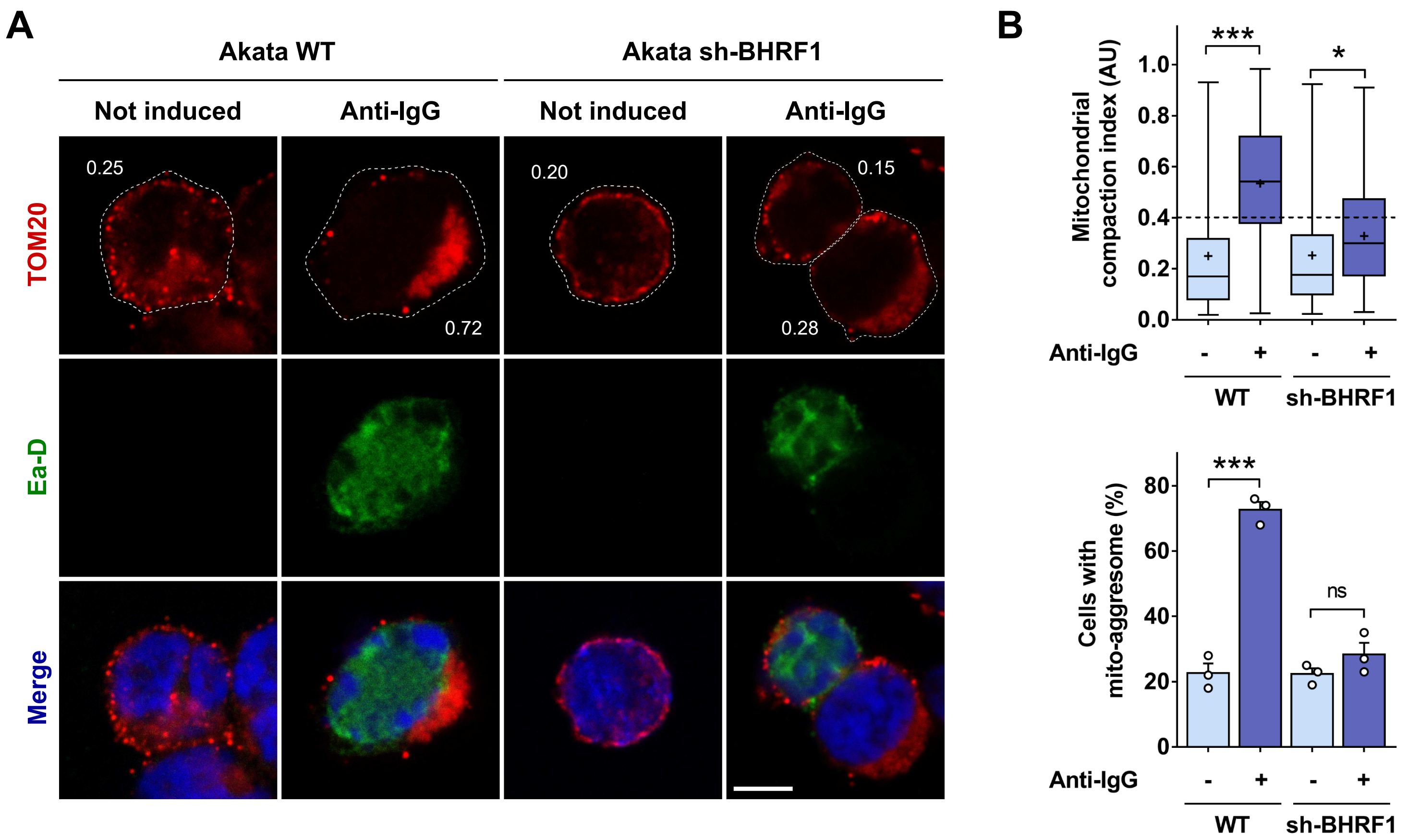


**Figure 1 – figure supplement 1. Knockdown of BHRF1 expression in Akata cells prevents mito-aggresome formation after EBV reactivation.** (**A-B**) Akata WT and sh-BHRF1 cells were treated or not with anti-human IgG for 24 h and then fixed. (**A**) Confocal images of cells immunostained for TOM20 and Ea-D. Nuclei were stained with DAPI. Values of mitochondrial CI are indicated. Scale bars: 5 µm. (**B**) Quantification of CI and percentage of cells with a mito-aggresome (n = 20 cells per condition).
Data represent the mean ± SEM of three independent experiments. ns = non-significant; * P < 0.05; *** P < 0.001 (Student's t-test).

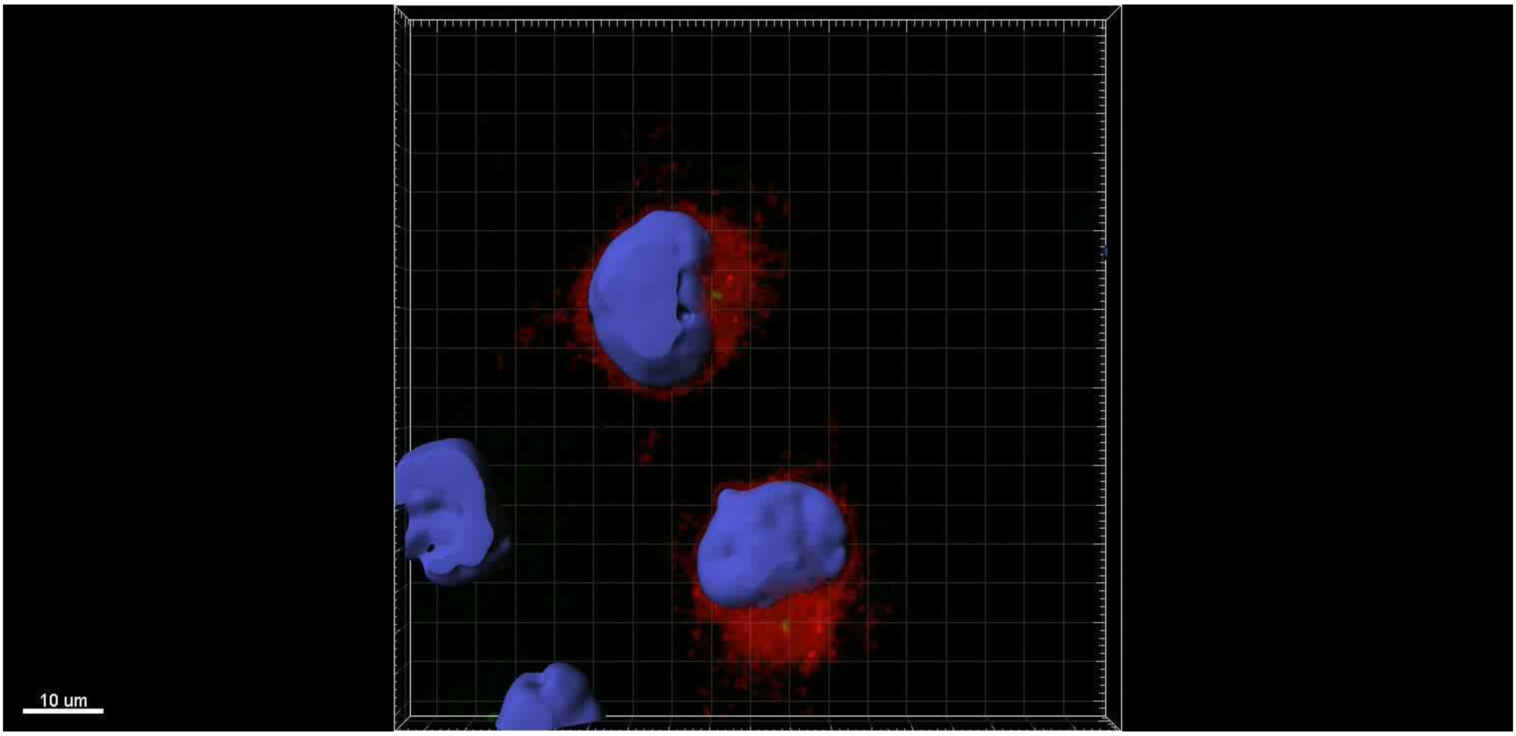


**Figure 1 – figure supplement 2. BHRF1 localizes in the proximity of the centrosome.** 3D Imaris surface reconstruction video of BHRF1-HA-transfected HeLa cells immunostained for pericentrin (centrosomal marker – green) and HA (BHRF1 – red). Nuclei were subsequently stained with DAPI. Scale bar: 10 µm.

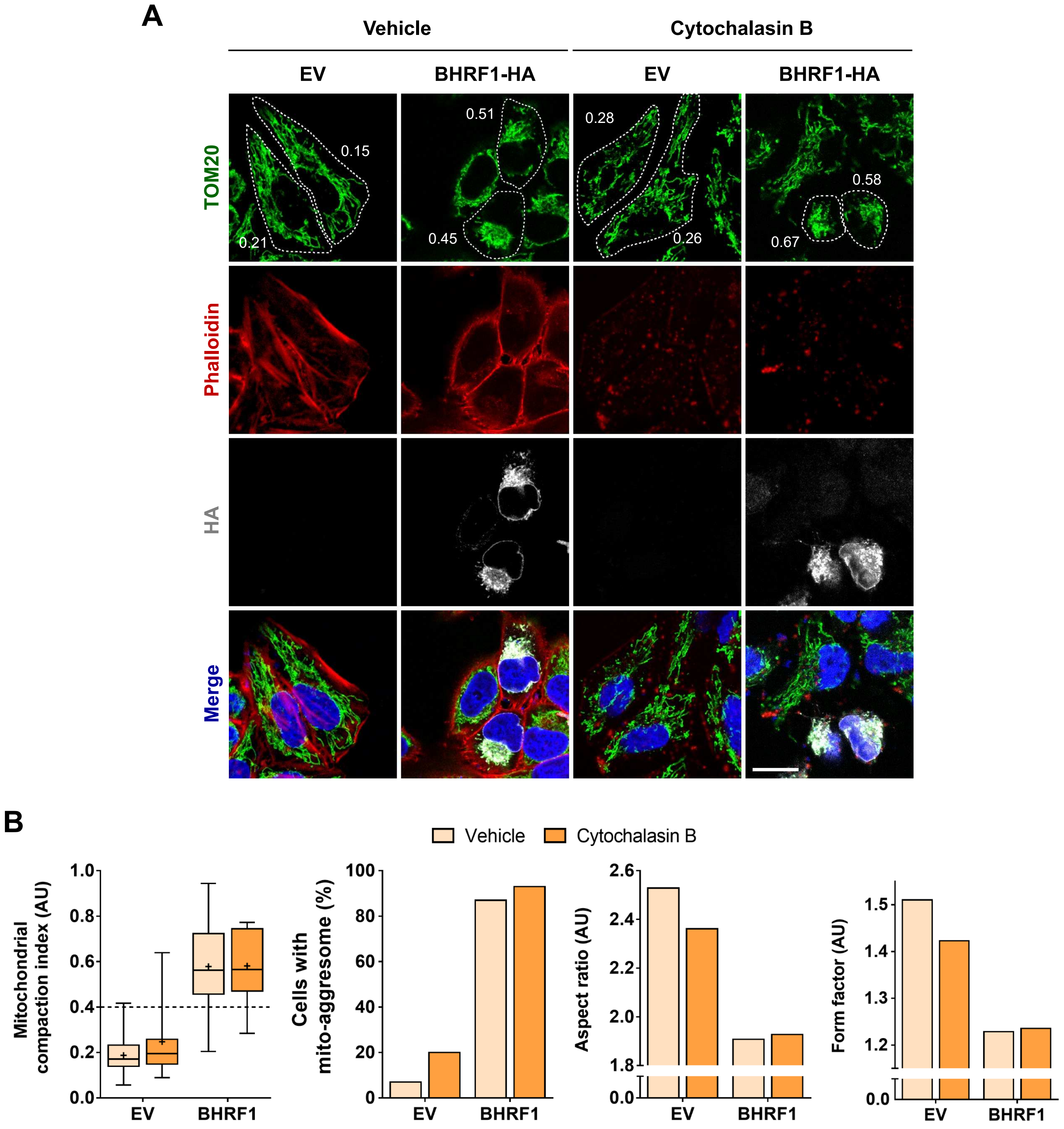


**Figure 1 – figure supplement 3. The actin network is not involved in the formation of BHRF1-induced mito-aggresomes.** (**A-B**) HeLa cells were transfected with BHRF1-HA plasmid (or EV) for 24 h. For the final 30 min, cells were treated with cytochalasin B to disrupt the F-actin network and then were fixed. (**A**) Confocal images. Cells were immunostained for TOM20 and HA. The F-actin network was labeled with phalloidin and the efficiency of the treatment with cytochalasin B was confirmed by the loss of F-actin staining. Nuclei were stained with DAPI. Values of mitochondrial CI are indicated on representative cells. Scale bar: 20 µm. (**B**) Quantification of CI, percentage of cells with a mito-aggresome and mitochondrial fission parameters (n = 20 cells per condition).

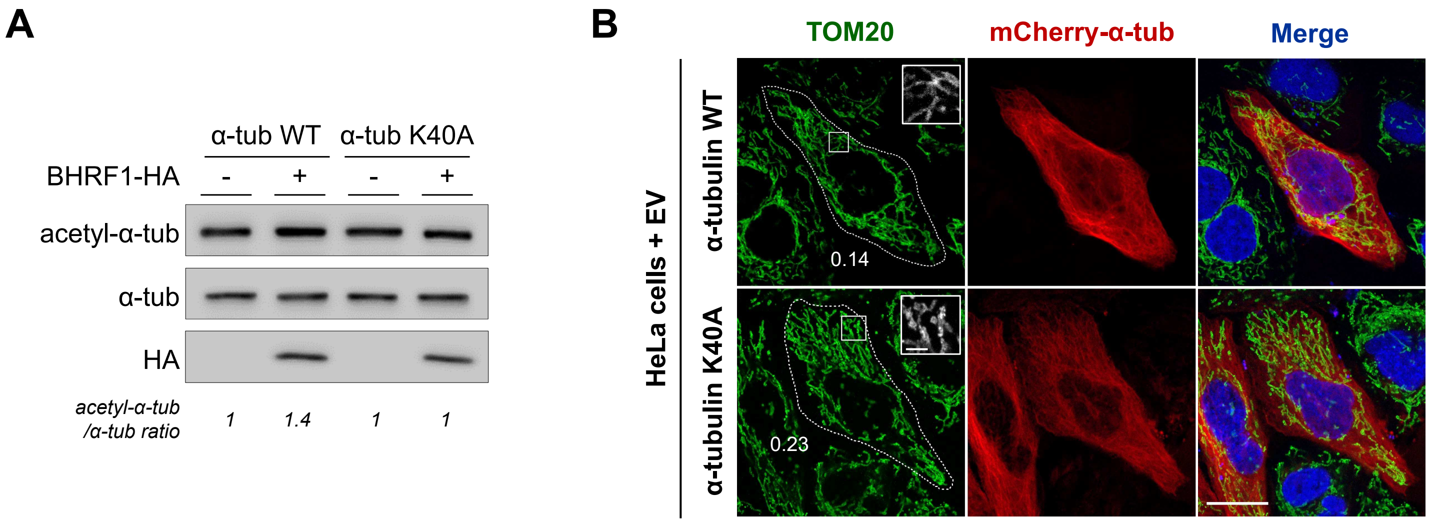


**Figure 3 – figure supplement 1. Impact of the non-acetylatable α-tubulin expression.** HeLa cells were co-transfected with plasmids encoding BHRF1-HA (or EV) and mCherry-α-tubulin K40A (or WT). (**A**) Immunoblot analysis of acetyl-α-tubulin, α-tubulin and HA. (**B**) Confocal images of EV-transfected cells immunostained for TOM20. Nuclei were stained with DAPI. Scale bars: 10 µm and 4 µm for insets. Values of mitochondrial CI are indicated on representative cells.

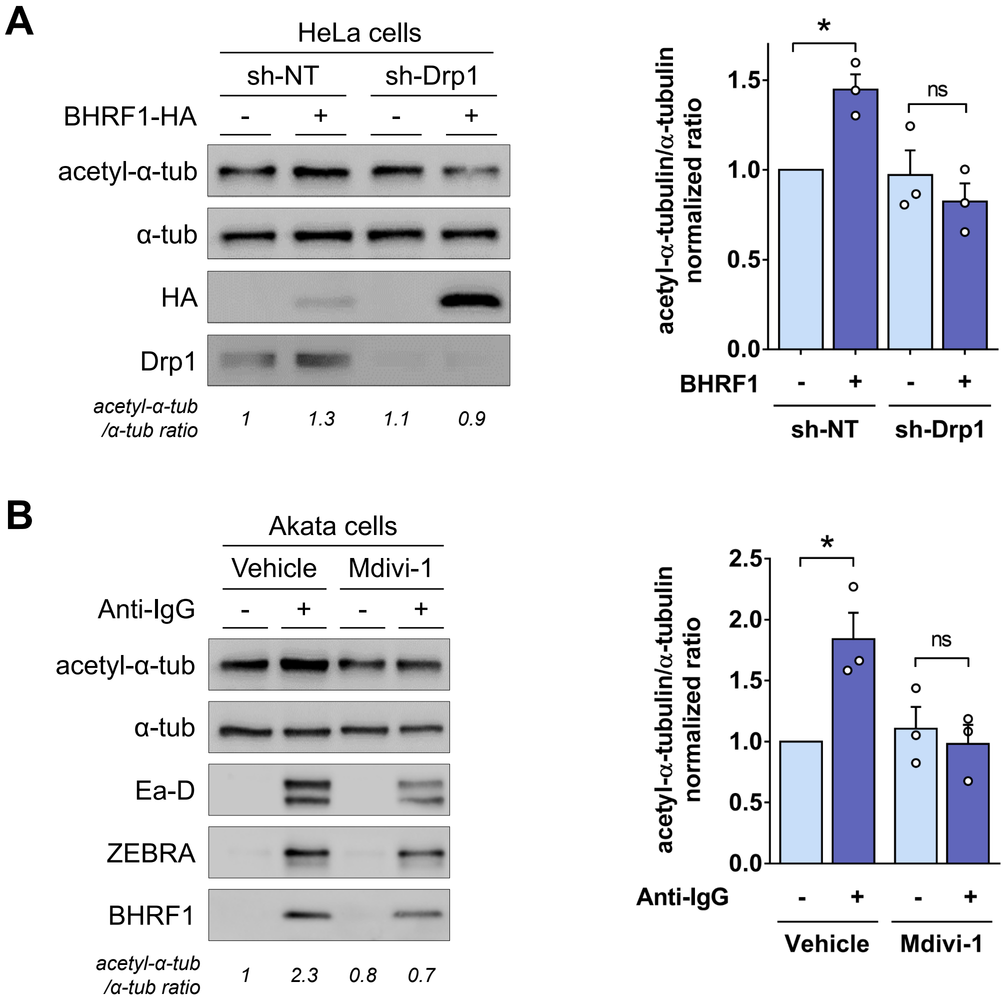


**Figure 3 – figure supplement 2. MT hyperacetylation required mitochondrial fission.** (**A**) HeLa cells expressing an sh-RNA against Drp1 (sh-Drp1) or a control sh-RNA (sh-NT) were transfected with BHRF1-HA for 24 h. *Left*, immunoblot analysis of acetyl-α-tubulin, α-tubulin, HA and Drp1. *Right*, normalized ratios of acetyl-α-tubulin to α-tubulin. (**B**) Akata cells are concomitantly treated for 24 h with anti-human IgG, to induce EBV reactivation, and with Mdivi-1, to inhibit Drp1. *Left*, immunoblot analysis of acetyl-α-tubulin, α-tubulin, Ea-D, ZEBRA and BHRF1. *Right*, normalized ratios of acetyl-α-tubulin to α-tubulin.

Data represent the mean ± SEM of three independent experiments. ns = non-significant; * $P < 0.05$ (Student's t-test).

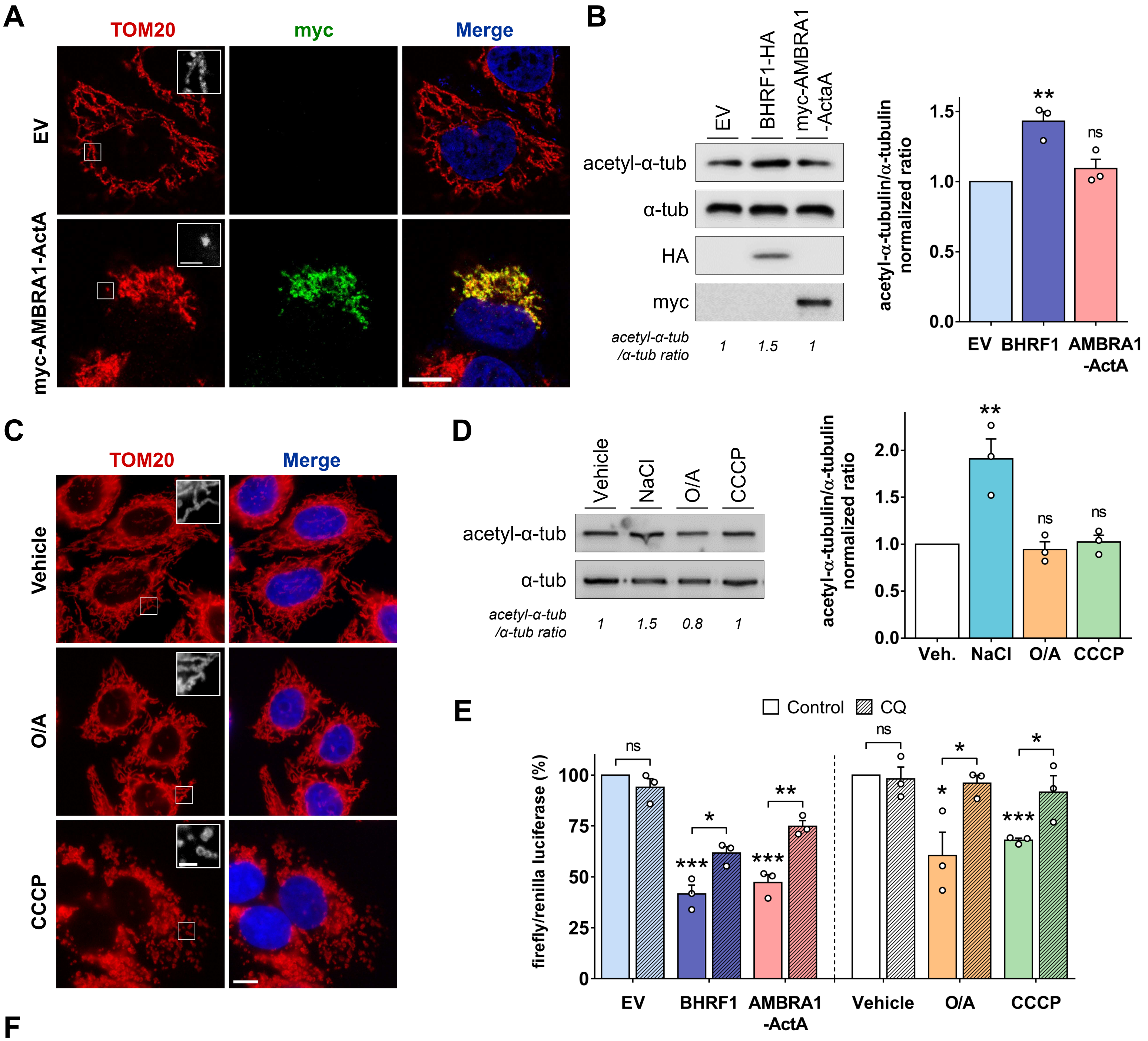


**F**

| Protein/Stress | Treatment | MT hyperacetylation | Mitochondrial fission | Mito-aggresome | Mitophagy induction | Inhibition of type I IFN |
|---|---|---|---|---|---|---|
| **BHRF1** | Transfected for 24h | Yes | Yes | Yes | Yes | Yes |
| **AMBRA1-ActA** | Transfected for 24h | No | Yes | Yes (*) (Strappazzon et al., 2015) | Yes (*) (Strappazzon et al., 2015) | Yes |
| **O/A** | 6 h [2.5 µM] / [800 nM] | No | No | No | Yes (*) (Lazarou et al., 2015) | Yes |
| **CCCP** | 4 h [10 µM] | No | Yes | No | Yes (*) (Narendra et al., 2008) | Yes |

**Figure 4 – figure supplement 1. Interplay between MT hyperacetylation and IFN inhibition in response to different mitophagy inducers.** (**A-B**) HeLa cells were transfected with plasmids encoding myc-AMBRA1-ActA or BHRF1-HA for 24 h. (**A**) Confocal images with insets (3X) of AMBRA1-ActA-transfected cells immunostained for TOM20 and myc. Nuclei were stained with DAPI. Scale bars: 10 µm and 2 µm for insets. (**B**) *Left*, immunoblot analysis of acetyl-α-tubulin, α-tubulin, HA and myc. *Right*, normalized ratios of acetyl-α-tubulin to α-tubulin. (**C-D**) HeLa cells were treated for 6 h with O/A or 4 h with CCCP to induce mitophagy. (**C**) Representative images with insets (3X) of cells immunostained for TOM20. Nuclei were stained with DAPI. Scale bars: 10 µm and 2 µm for insets. (**D**) *Left*, immunoblot analysis of acetyl-α-tubulin and α-tubulin. Treatment with NaCl (30 min) was used as a positive control of hyperacetylation. *Right*, normalized ratios of acetyl-α-tubulin to α-tubulin. (**E**) Luciferase reporter assay on HEK293T cells treated with various mitophagy inducers. Cells are either transfected with BHRF1 or AMBRA1-ActA, or treated with O/A or CCCP. Before lysis, cells are treated or not with CQ for 4 h. Activation of the IFN-β promoter was analyzed 24 h post-transfection, or after indicated treatment. Firefly/renilla luciferase ratios were calculated and normalized to control conditions (EV or vehicle). (**F**) Summary table of the impact of mitophagy inducers on MT hyperacetylation and IFN inhibition. Results indicated by a star have been previously reported.
Data represent the mean ± SEM of three independent experiments. ns = non-significant; * P < 0.05; ** P < 0.01; *** P < 0.001 (Student's t-test).

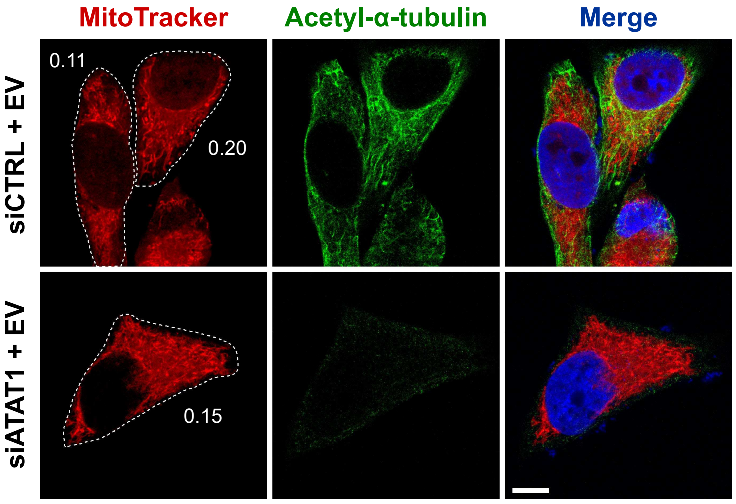


**Figure 5 – figure supplement 1. Loss of ATAT1 has no impact on mitochondrial network.** After knockdown of ATAT1, HeLa cells were transfected with an EV and fixed 24 h post-transfection. Mitochondria were labeled with MitoTracker, and cells were immunostained for acetyl-α-tubulin. Nuclei were stained with DAPI. Values of mitochondrial CI are indicated on representative cells. Scale bar: 20 µm.

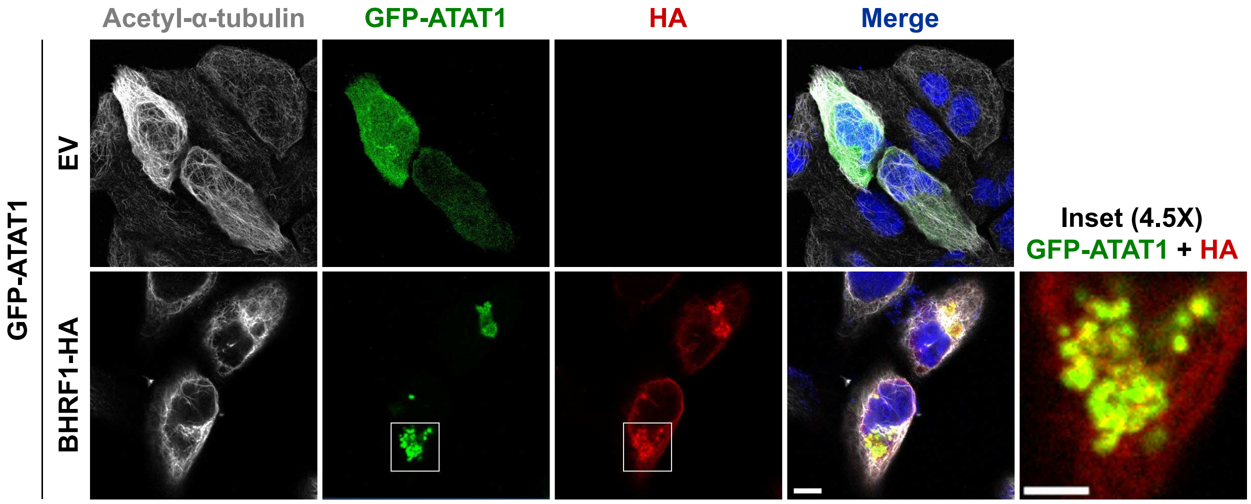


**Figure 5 – figure supplement 2. ATAT1 colocalizes with BHRF1.** HeLa cells were co-transfected with GFP-ATAT1 and BHRF1-HA (or EV) for 24 h. Confocal images of cells immunostained for HA and acetyl-α-tubulin. Nuclei were stained with DAPI. Inset shows colocalization between ATAT1 and BHRF1. Scale bars: 10 µm and 5 µm for inset.

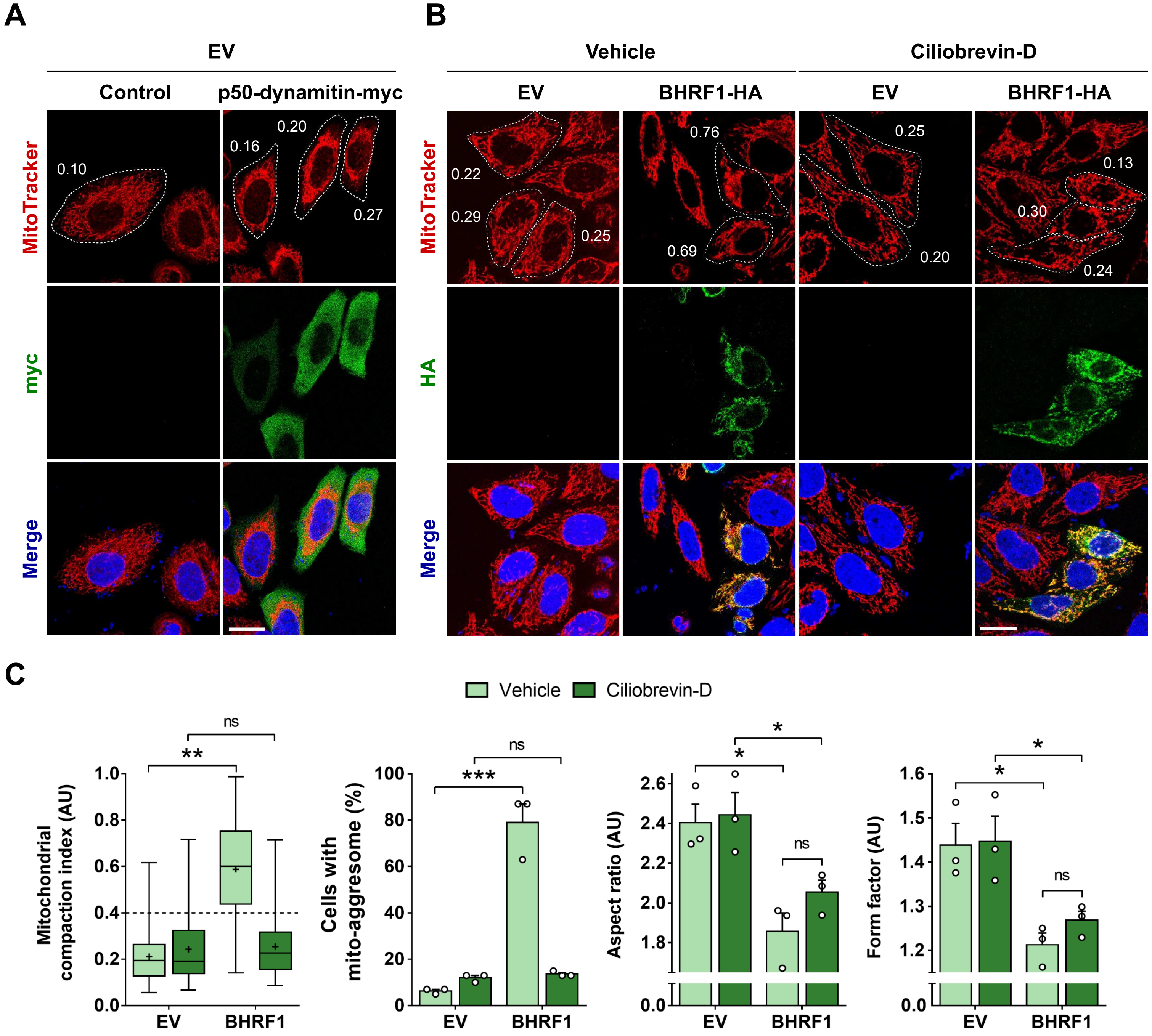


**Figure 6 – figure supplement 1. Dynein inhibition prevents BHRF1-induced mito-aggresomes. (A)** Expression of p50-dynamitin-myc has no impact on the mitochondrial network. Confocal images of HeLa cells co-transfected with plasmids encoding EV and p50-dynamitin-myc for 24 h. Mitochondria were labeled with MitoTracker, and cells were immunostained for a myc-tag. Nuclei were stained with DAPI. Values of mitochondrial CI are indicated on representative cells. Scale bar: 20 µm. (**B-C**) HeLa cells were transfected for 24 h with BHRF1-HA (or EV) and treated overnight with ciliobrevin-D. (**B**) Confocal images. Mitochondria were labeled with MitoTracker, cells immunostained for HA and nuclei stained with DAPI. Values of mitochondrial CI are indicated on representative cells. Scale bar: 20 µm. (**C**) Quantification of CI, percentage of cells with a mito-aggresome and mitochondrial fission parameters (n = 20 cells per condition).
Data represent the mean ± SEM of three independent experiments. ns = non-significant; * $P < 0.05$; ** $P < 0.01$; *** $P < 0.001$ (Student's t-test).

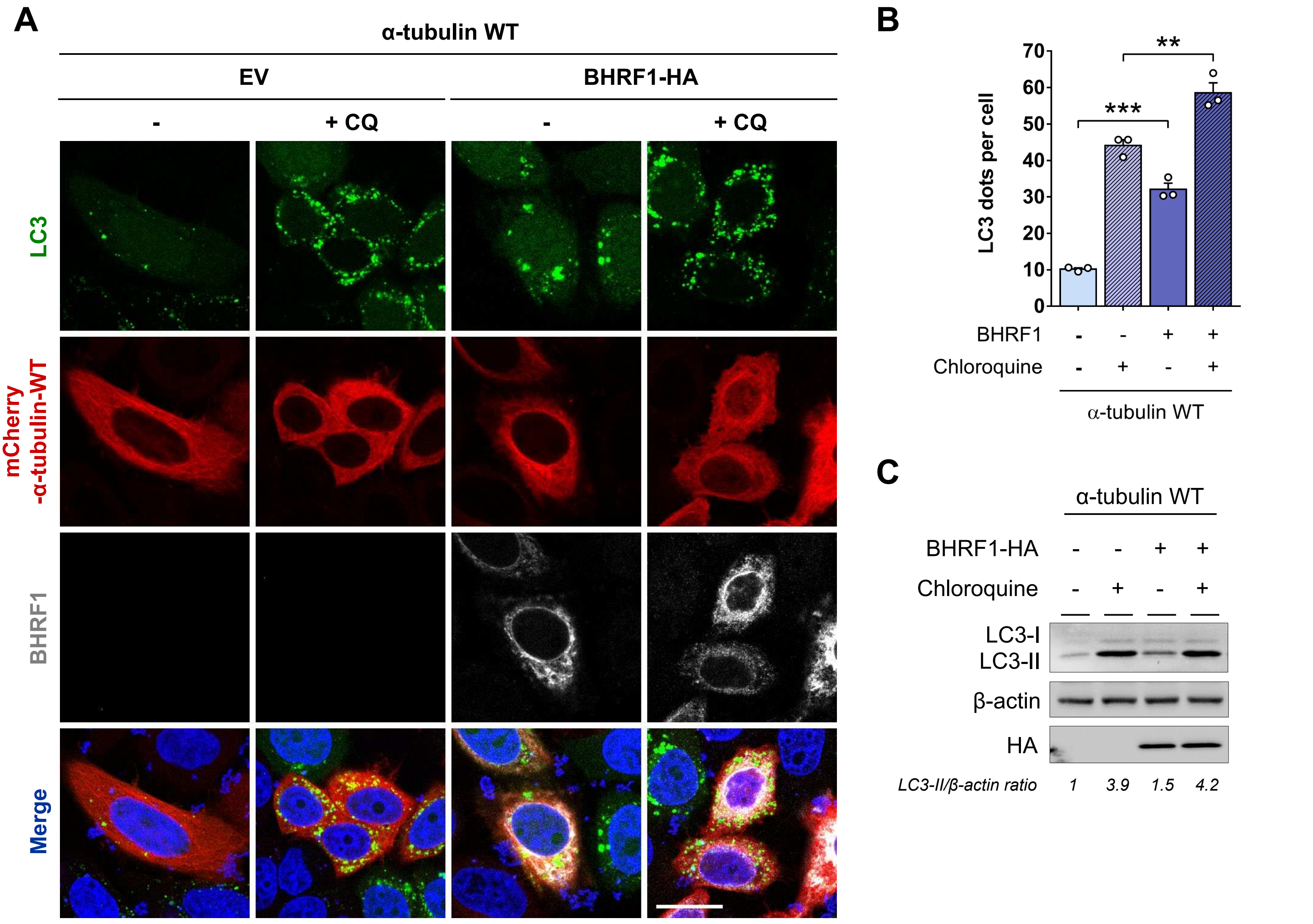


**Figure 7 – figure supplement 1. BHRF1 stimulates autophagy when α-tubulin WT is expressed.** (**A-C**) HeLa cells co-transfected for 24 h with plasmids encoding BHRF1-HA (or EV) and mCherry-α-tubulin WT. Cells were treated with CQ when indicated. (**A**) Confocal images. Cells were immunostained for BHRF1 and LC3 and nuclei were stained with DAPI. Scale bar: 20 µm. (**B**) Quantification of LC3 dots (n = 30 cells per condition). (**C**) Immunoblot analysis of LC3 and BHRF1-HA expression. β-actin was used as a loading control.
Data represent the mean ± SEM of three independent experiments. ** P < 0.01; *** P < 0.001 (Student's t-test).

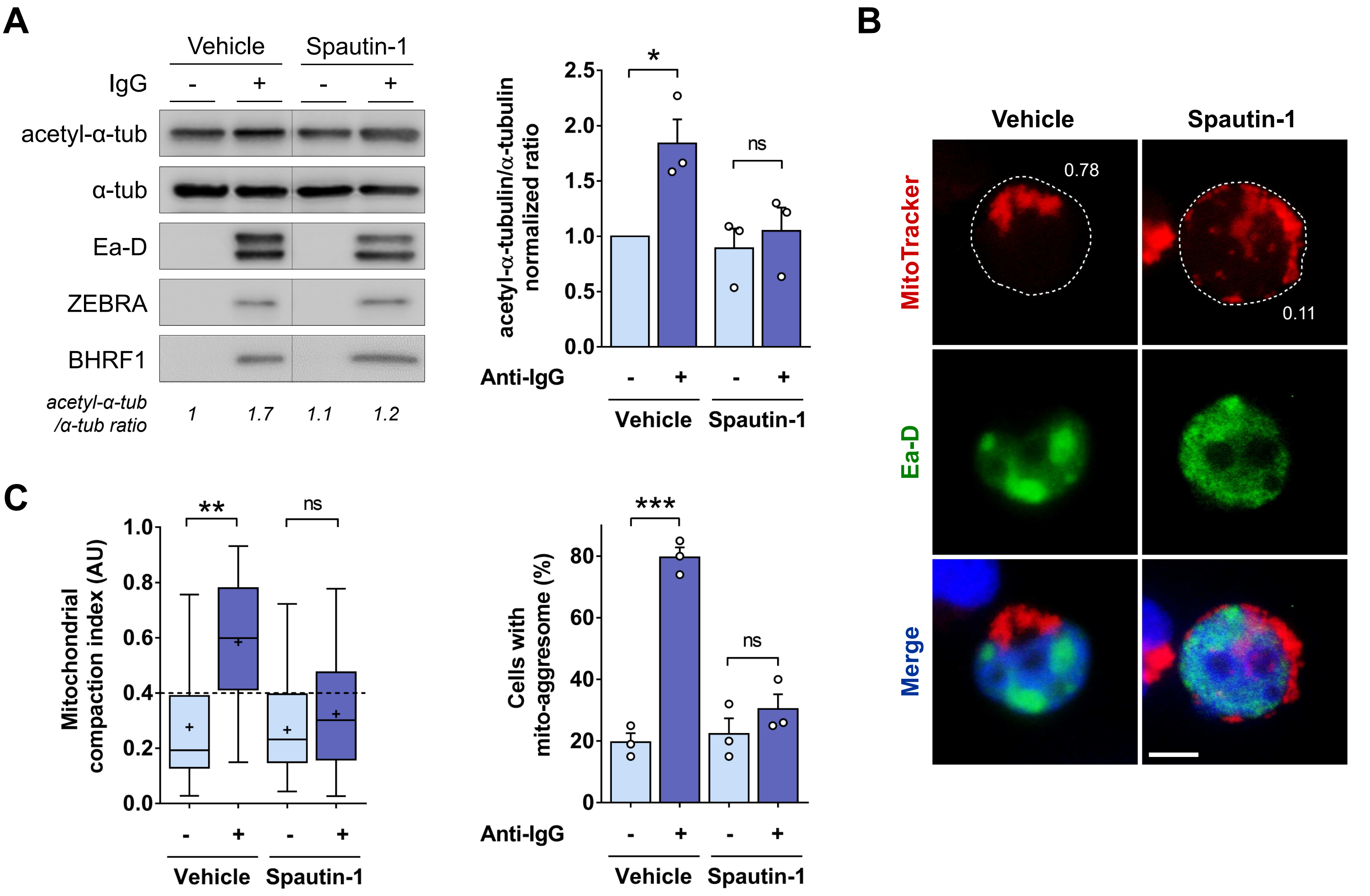


**Figure 8 – figure supplement 1. Autophagy inhibition prevents MT hyperacetylation and mito-aggresome formation during EBV reactivation.** (**A-C**) EBV reactivation was induced in Akata cells by treatment with anti-human IgG for 24 h. At the same time, cells were treated with spautin-1 (or vehicle) for 24 h to block autophagy. (**A**) *Left*, immunoblot analysis of acetyl-α-tubulin, α-tubulin, Ea-D, ZEBRA and BHRF1. *Right*, normalized ratios of acetyl-α-tubulin to α-tubulin. (**B**) Representative images where mitochondria were labeled with MitoTracker, cells immunostained for Ea-D and nuclei stained with DAPI. Values of mitochondrial CI are indicated on each cell. Scale bar: 10 µm. (**C**) Quantification of CI and percentage of cells with a mito-aggresome (n = 20 cells per condition).

Data represent the mean ± SEM of three independent experiments. ns = non-significant; * P < 0.05; ** P < 0.01; *** P < 0.001 (Student's t-test).
